# Integrated transcriptomic analysis of human induced pluripotent stem cell-derived osteogenic differentiation reveals a regulatory role of KLF16

**DOI:** 10.1101/2024.02.11.579844

**Authors:** Ying Ru, Meng Ma, Xianxiao Zhou, Divya Kriti, Ninette Cohen, Sunita D’Souza, Christoph Schaniel, Susan M. Motch Perrine, Sharon Kuo, Oksana Pichurin, Dalila Pinto, Genevieve Housman, Greg Holmes, Eric Schadt, Harm van Bakel, Bin Zhang, Ethylin Wang Jabs, Meng Wu

## Abstract

Osteogenic differentiation is essential for bone development, metabolism, and repair; however, the underlying regulatory relationships among genes remain poorly understood. To elucidate the transcriptomic changes and identify novel regulatory genes involved in osteogenic differentiation, we differentiated mesenchymal stem cells (MSCs) derived from 20 human iPSC lines into preosteoblasts (preOBs) and osteoblasts (OBs). We then performed transcriptome profiling of MSCs, preOBs and OBs. The iPSC-derived MSCs and OBs showed similar transcriptome profiles to those of primary human MSCs and OBs, respectively. Differential gene expression analysis revealed global changes in the transcriptomes from MSCs to preOBs, and then to OBs, including the differential expression of 840 genes encoding transcription factors (TFs). TF regulatory network analysis uncovered a network comprising 451 TFs, organized into five interactive modules. Multiscale embedded gene co-expression network analysis (MEGENA) identified gene co-expression modules and key network regulators (KNRs). From these analyses, *KLF16* emerged as an important TF in osteogenic differentiation. We demonstrate that overexpression of *Klf16* in vitro inhibited osteogenic differentiation and mineralization, while *Klf16^+/-^* mice exhibited increased bone mineral density, trabecular number, and cortical bone area. Our study underscores the complexity of osteogenic differentiation and identifies novel regulatory genes such as *KLF16*, which plays an inhibitory role in osteogenic differentiation both in vitro and in vivo.

## Introduction

Mesenchymal stem cells (MSCs) are multipotent cells that can differentiate into a variety of lineages, including osteoblasts (OBs), chondrocytes, and adipocytes (Pittenger et al., 1999). The multilineage potential, self-renewal capacity, and immune-modulation functions of MSCs have made them a promising therapeutic tool for cell therapies and regenerative medicine (Bruder et al., 1994; Han et al., 2019). Osteogenic differentiation from MSCs plays a pivotal role in bone development, homeostasis, and repair. This process replenishes the pool of osteoblasts and ensures that there are enough cells available for bone formation and repair. It is highly regulated by multiple mechanisms and previous research has highlighted the roles of various genes, including those coding for transcription factors (TFs) and signaling pathways such as TGF-β/BMP, Notch, Hedgehog, Wnt, Hippo, and estrogen receptor signaling (Thomas and Jaganathan, 2022). Precise and robust transcriptional regulation required during osteogenic differentiation is achieved by networks of transcriptional regulators. For example, *RUNX2* is a master transcription factor in early osteogenic differentiation (Komori et al., 1997; Schroeder et al., 2005). TFs *SP7, ATF4, TEAD4*, and *KLF4* work alongside *RUNX2* to regulate osteogenesis (Nakashima et al., 2002; Suo et al., 2020; Yang et al., 2004; Yu et al., 2021). The broader TF interplay in osteogenic differentiation remains largely uncharted, even with insights from ENCODE (Encyclopedia of DNA Elements) data that reveal the co-association of human TFs in a combinatorial and context-specific manner (Gerstein et al., 2012), revealing a need for further investigation.

Transcriptome profiling can uncover the key genes essential for bone formation and growth, offering valuable insights into bone development and the underlying mechanisms of bone-related diseases. However, due to the invasiveness required to obtain human primary MSCs and the suboptimal conditions for preserving available samples, large-scale human transcriptomics projects, such as GTEx, do not include data from bone and cartilage. Furthermore, studies focusing on these tissues are often limited to disease contexts. Induced pluripotent stem cell (iPSC)-derived systems, with their pluripotency and differentiation capabilities, offer an avenue for studying skeletal development. iPSC-derived sclerotome models have been utilized to identify gene expression signatures and regulatory mechanisms crucial to key developmental stages of endochondral ossification (Lamande et al., 2023; Nakajima et al., 2018; Tani et al., 2023). Previous studies for osteogenic differentiation from iPSC-derived MSCs included only one to two MSC cell lines (Matsuda et al., 2020; Rauch et al., 2019), limiting their power for comprehensive transcriptomic analyses. In this study, 20 human iPSC lines, each from a healthy individual, were used to investigate osteogenic differentiation of MSCs by RNA sequencing (RNA-seq) at three key stages: MSC, preosteoblast (preOB), and osteoblast (OB). Our findings highlight dynamic gene expression changes during osteogenic differentiation and identify the regulatory role of TF KLF16 as an inhibitor of osteogenic differentiation. This work provides insights into osteogenic differentiation and sheds light on novel therapeutic targets for bone diseases.

## Results

### Transcriptome profiles of human iPSC-derived MSCs and OBs are similar to those of primary MSCs and OBs

Human iPSCs were generated from peripheral blood mononuclear cells (PBMCs) or skin fibroblasts of 20 healthy individuals (9 males and 11 females) and differentiated into MSC, preOB, and OB stages (Figure 1A; Supplementary Figure 1A, 1B, and 1H; Supplementary Table 1). To abrogate the epigenetic memory from their source cell types and minimize in vitro transcriptional and differentiation heterogeneity, the established iPSC lines were cultured for at least 16 passages and underwent quality control (Polo et al., 2010). The iPSCs were then differentiated into MSCs and sorted for the CD105+/CD45- cell population (Giuliani et al., 2011; Kang et al., 2015) (Supplementary Figure 1C). No significant differences were found in the percentage of this cell population between PBMC and fibroblast origins (Supplementary Figure 1D). After two weeks of expansion, the iPSC-derived MSCs were confirmed to exhibit mesenchymal characteristics, evidenced by strong expression of positive MSC surface markers (CD29, CD73, CD90, and CD105) and minimal expression of negative markers (CD31, CD34, and CD45) at both the protein level by flow cytometry (Supplementary Figure 1E and 1F) and mRNA level by RNA-seq (Supplementary Figure 1G). The MSCs were cultured in osteogenic differentiation medium for 7 days to generate preOBs and for 21 days to produce OBs. After 21 days of osteogenic differentiation, the OBs showed alkaline phosphatase (ALP) enzyme activity and mineralization, indicating the state of functional OBs (Figure 1B).

**Figure 1.**
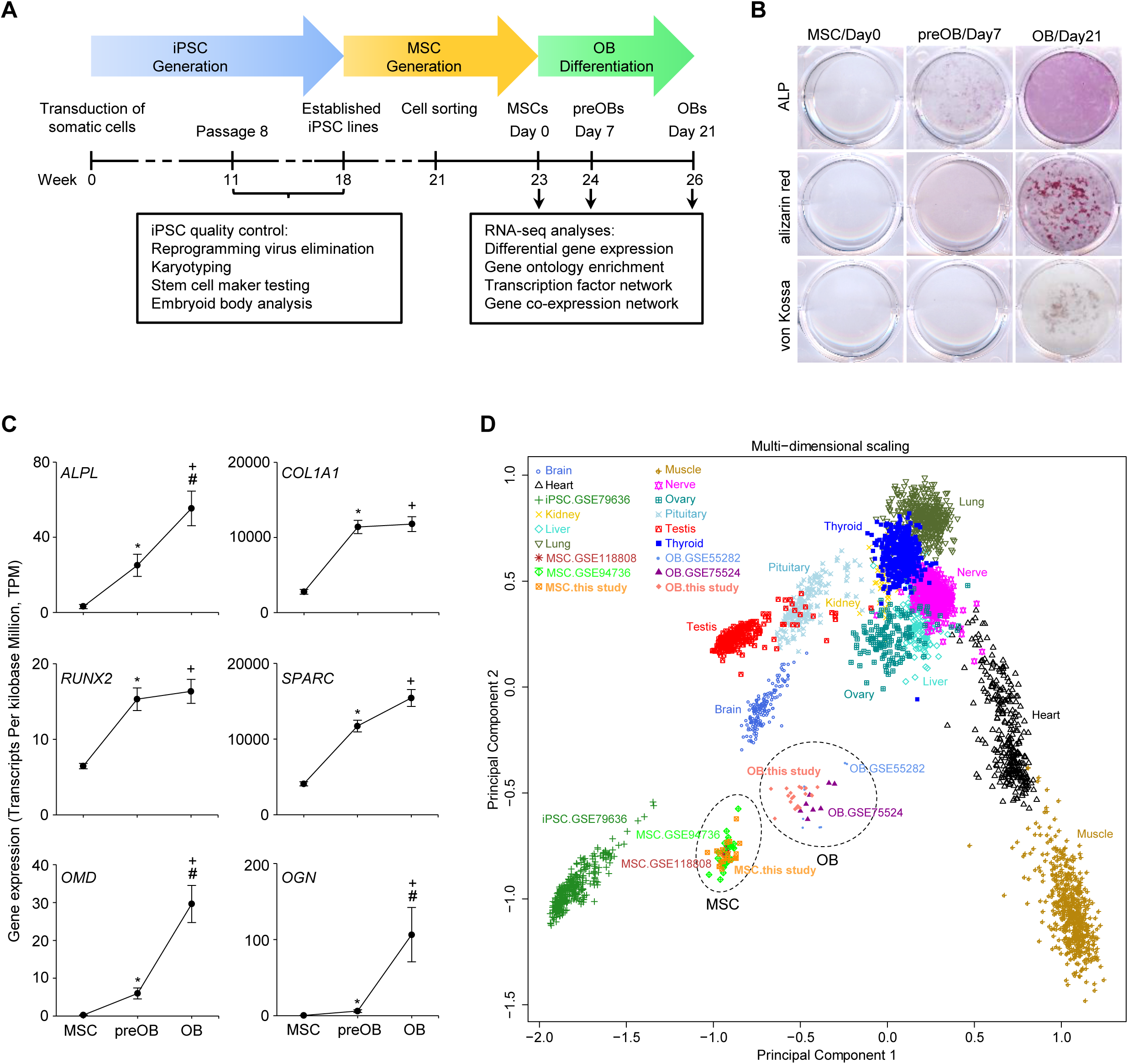
Generation of healthy human iPSCs and osteogenic differentiation transcriptomic data. **(A)** Flowchart of iPSC establishment, iPSC-derived MSC generation, MSC to OB differentiation (preOBs, preosteoblasts; OBs, osteoblasts), and RNA-seq data generation and analyses. **(B)** In vitro osteogenic differentiation of MSCs (Day 0), preOBs (Day 7), and OBs (Day 21) stained with alkaline phosphatase (ALP), alizarin red, and von Kossa. **(C)** Expression level of known osteogenic genes at the three osteogenic stages. Data are shown as mean ± SEM. * MSC vs preOB, # preOB vs OB, or + MSC vs OB, adjusted p value < 0.0001. **(D)** Principal component analysis (PCA) of RNA-seq data from our iPSC-derived MSCs and differentiated OBs as well as previously published human primary MSCs and OBs, iPSCs, and other tissues in GTEx.

MSCs, preOBs, and OBs derived from the 20 individuals were harvested and a total of 60 RNA-seq libraries were generated (Figure 1A; Supplementary Table 1). We initially employed bulk RNA-seq rather than single-cell RNA-seq (scRNA-seq) because the latter recovers only sparse transcriptional data from individual cells, thus limiting comprehensive expression analysis and the development of robust prediction models for regulatory and gene co-expression networks (Li and Wang, 2021). RNA-seq data revealed increased expression of OB lineage markers (*ALPL, COL1A1, RUNX2, SPARC, OMD,* and *OGN*) with characteristic temporal expression patterns during osteogenic differentiation (Figure 1C). We further compared our MSC and OB RNA-seq datasets with previously published human primary MSC and OB datasets, human iPSC datasets, and GTEx datasets for different tissues derived from the three germ layers (ectoderm, endoderm, and mesoderm) and germ cells (Al-Rekabi et al., 2016; Ardlie et al., 2015; Ma et al., 2019; Roforth et al., 2015; Rojas-Peña et al., 2014). By principal component analysis (PCA), our MSC and OB datasets clustered with those of published primary MSCs and OBs, respectively, and were discrete from iPSCs and other tissue datasets, further validating the cell identities of our MSCs and OBs (Figure 1D).

### Differential transcription profiles of human iPSC-derived MSCs, preOBs, and OBs

During in vitro osteogenic differentiation, we detected the expression of 17,795 unique genes across all stages, of which 79% were protein-coding and 21% were noncoding (Figure 2A). Seventy percent or 9,724 of the protein-coding genes were differentially expressed (DE) between the osteogenic stages (MSC to preOB, or preOB to OB) (fold change ≥ 1.2 and adjusted p value ≤ 0.05). Sixty percent (2,297) of the noncoding genes also were DE (Figure 2A), of which 2,249 (98% of DE noncoding) were long noncoding RNA (lncRNA) genes. By intersecting with the human TF repertoire including over 1,600 known or likely human TFs (Lambert et al., 2018), we found that nine percent (840 genes) of the DE coding genes were TFs (Figure 2B). A chi-square test revealed a significant correlation between TFs and differentially expressed genes (DEGs) (X^2^ = 27.4, p value < 0.0001). There were 5,715 up-regulated genes and 4,409 down-regulated genes from the MSC to the preOB stage, and 3,342 up-regulated genes and 2,835 down-regulated genes from the preOB to the OB stage, revealing that the majority of the DEGs were up-regulated during osteogenic differentiation (Figure 2C-E; Supplementary Table 2). Among the top statistically significant DEGs were known osteogenesis-associated genes, including a SMAD ubiquitination regulatory factor *SMURF2* (Kushioka et al., 2020), natriuretic peptide precursor B *NPPB* (Aza-Carmona et al., 2014), hypoxia-inducible factor 3 subunit alpha *HIF3A* (Zhu et al., 2014), a TF *ZBTB16* (Onizuka et al., 2016), a Wnt signaling pathway family member *WNT7B* (Yu et al., 2020), osteoinductive factor *OGN* (Kukita et al., 1990), a SFRP family (modulators of Wnt signaling) member *SFRP2* (Yang et al., 2020), and a glycoprotein member of the glycosyl hydrolase 18 family *CHI3L* (Chen et al., 2017). There were also many genes with no defined function in bone, including those that play critical roles in RNA binding activity (*EBNA1BP2*), maintenance of protein homeostasis (*PSMD2*), arachidonic acid metabolism (*PRXL2B*), hemostasis and antimicrobial host defense (*FGB*), embryonic and induced pluripotent stem cells (*PCSK2*), as well as cell proliferation, differentiation, and migration (*FGFBP1*). A greater number of DEGs with larger changes in gene expression were identified between the MSC and preOB stages compared to the changes between the preOB and OB stages, even though the differentiation interval was half as long between the MSC and preOB stages (one week) as it was between the preOB and OB stages (two weeks) (Figure 2C-E).

**Figure 2.**
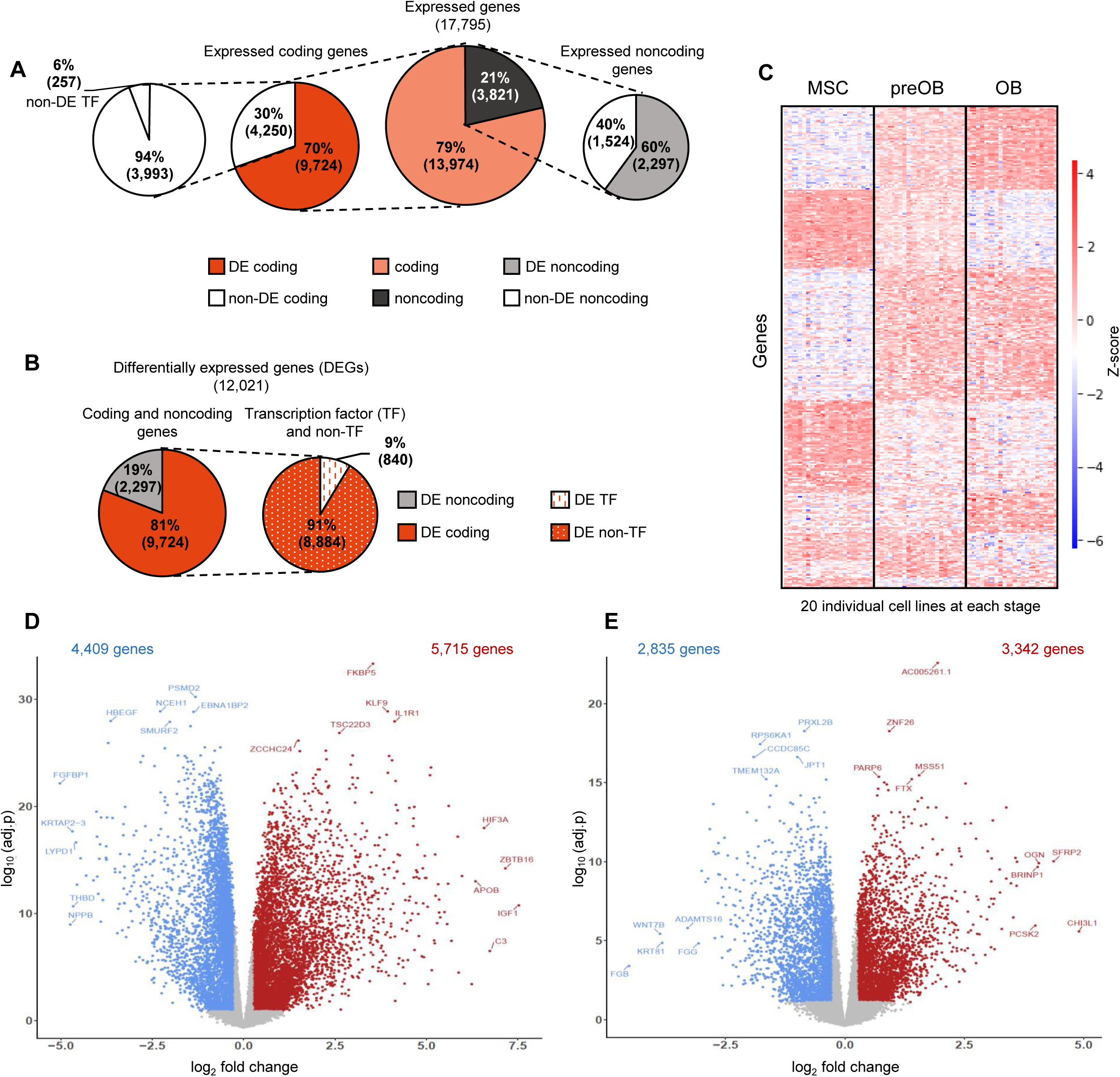
Gene expression profile during osteogenic differentiation. **(A)** Gene expression during osteogenic differentiation with the total number of expressed genes and percentages of expressed coding and noncoding genes (middle). Number and percentages of differentially expressed (DE) and non-differentially expressed (non-DE) coding genes (left; TFs and non-TFs (white pie chart)) and of noncoding genes (right). **(B)** DEGs during osteogenic differentiation with total and percentages of coding and noncoding genes (left), and TFs and non-TFs (right). **(C)** Heatmap showing hierarchical clustering of 60 RNA-seq datasets from 20 iPSC-derived MSC, preOB, and OB lines (columns) and significant differentially expressed genes (DEGs) (rows), fold change ≥ 1.2 and adjusted p value < 0.05. Up-regulated and down-regulated gene expression is colored in red and blue, respectively. **(D)** and **(E)** Volcano plots illustrate the distribution of down- and up-regulated genes (blue and red, respectively) with adjusted p values and fold changes when comparing gene differential expression from MSC to preOB in **(D)** and preOB to OB stages in **(E)**. Cutoffs of fold change ≥ 1.2 and adjusted p value < 0.05 were applied to define DEGs. The total number of up-regulated and down-regulated genes are noted at the top (red and blue, respectively). The genes are labeled for the top five up-regulated (red, bottom right), downregulated (blue, bottom left), and statistically significant down-regulated (blue, top left) and up-regulated (red, top right).

### Transcription factor regulatory network during human osteogenic differentiation

By PCA, we found that the DEG expression profiles segregated all of the samples by their differentiation stage, with MSCs being most distinct from preOBs and OBs, which is consistent with the larger number of DEGs present between the MSC and preOB stages than the later stages (Figure 3A). Of note, when we used only the 840 differentially expressed TFs, their expression profiles also segregated the three osteogenic stages (Figure 3B).

**Figure 3.**
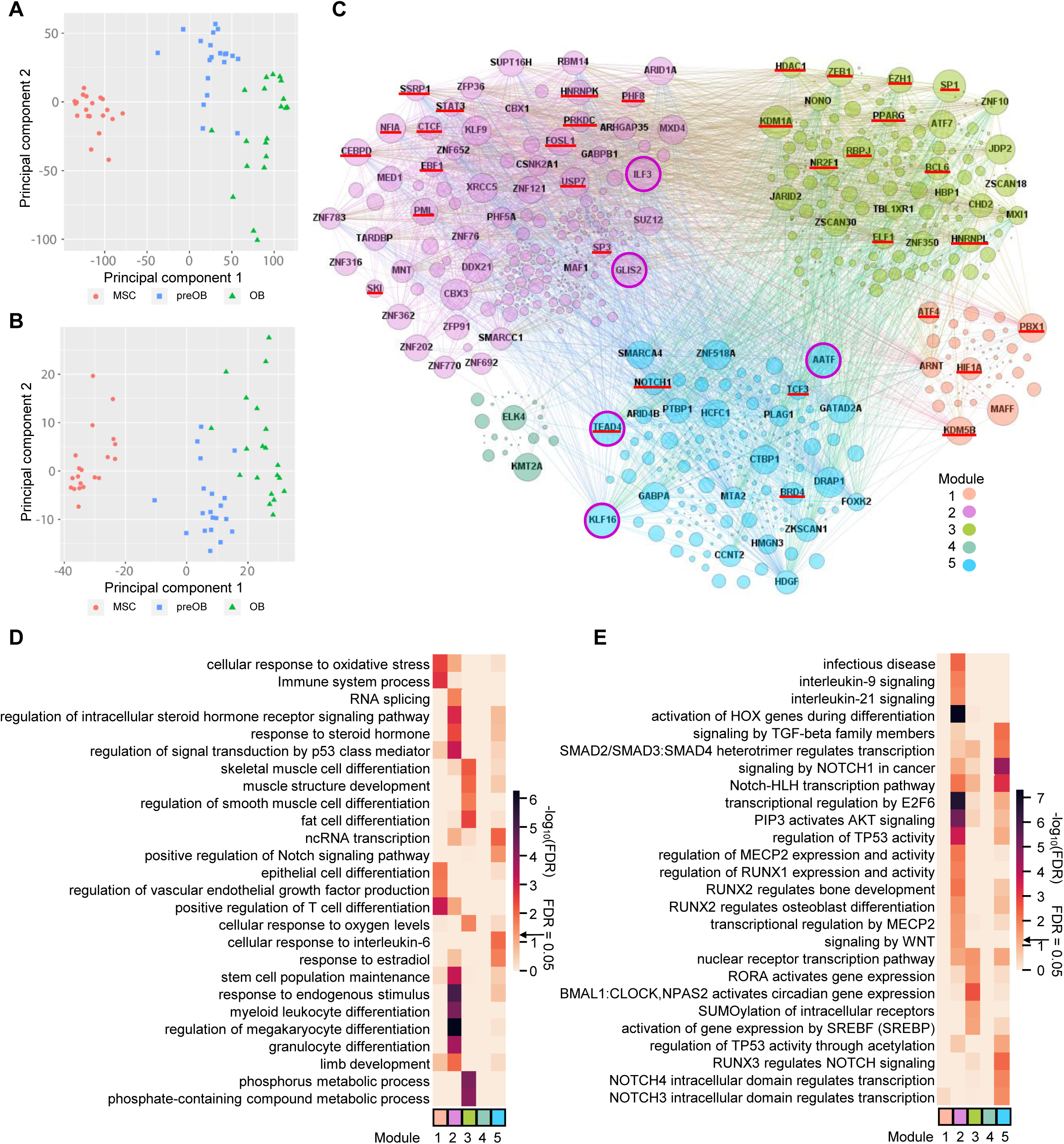
TF regulatory network in osteogenic differentiation. **(A)** Principal component analysis (PCA) of all 20 healthy cell lines at three stages of osteogenic differentiation using all differentially expressed genes (DEGs). **(B)** PCA using only differentially expressed TF genes. **(C)** TF regulatory network during osteogenic differentiation. Each node represents a TF, with known bone formation associated regulators underlined in red. Two nodes are connected by a line where ReMap data suggest regulation and our RNA-seq data suggest the association between them. Nodes labeled with the gene name represent the top 100 strongest TFs based on betweenness centrality. The size of the nodes reflects the regulation strength of the TF, with the top 5 strongest circled in pink. **(D)** and **(E)** Top significantly enriched GO BP terms in **(D)** and Reactome pathways in **(E)** of TFs in each network module.

TFs regulate not only the transcription of other protein-coding genes and noncoding RNA genes but also their own expression. We focused on the TF biological cooperativity with respect to TF-TF regulation, which has not been extensively investigated in osteogenic differentiation due to the limited availability of DNA-binding data from MSCs, preOBs, and OBs. To predict the TF-TF regulatory network during osteogenic differentiation, we constructed a network using our RNA-seq data and the database ReMap, which integrated and analyzed 5,798 human ChIP-seq and ChIP-exo datasets from public sources providing a transcriptional regulatory repertoire to predict target genes (Chèneby et al., 2020). ReMap covers 1,135 transcriptional regulators, including primarily TFs as well as coactivators, corepressors, and chromatin-remodeling factors, with a catalog of 165 million binding peaks. We filtered out TFs that lacked associations in ReMap, as they are less likely to have direct binding activities in regulating the transcription of TF genes. To further assure the reliability and accuracy of the regulatory relationships in our osteogenic differentiation datasets, we then assessed regulations among them by computing correlation coefficients of gene expression, resulting in 451 TFs in the network. After the application of a partitioning algorithm, the network was organized into five interconnected modules (Bastian et al., 2009) (Figure 3C; Supplementary Table 3). The network showed that TFs potentially regulate other TFs both internal and external to their respective modules, revealing a highly complex network of transcriptional regulators. We identified the top 100 TFs determined by their betweenness centrality, which coincided with their power to regulate others in the network (Bastian et al., 2009) (Figure 3C; Supplementary Table 3). Among them, TF genes known to regulate osteogenic differentiation were present in different modules, such as *ATF4* (Yang et al., 2004) in Module 1, *FOSL1* (Krum et al., 2010) in Module 2, *ZEB1* (Fu et al., 2020) in Module 3, and *TEAD4* (Suo et al., 2020) in Module 5 (Supplementary Table 3). Of interest, TF *KLF16* in Module 5, which previously was not demonstrated to be involved in bone formation and development, was identified as a key player in the regulatory network. It ranks as the fifth top regulator, directly interacting with the second top regulator *TEAD4*, and shows a high degree of association with 315 TFs (Supplementary Table 3).

Gene ontology (GO) and Reactome pathway (RP) analyses revealed regulatory functions and pathways specific to each module (FDR < 0.05; Figure 3D and 3E; Supplementary Table 3). As examples for the GO biological process (BP) terms, Module 3 was enriched for skeletal muscle cell differentiation and muscle structure development, associated with 15 TF genes (*ZBTB18*, *MEF2A*, *EGR1*, *TCF7L2*, *EGR2*, *EPAS1*, *SRF*, *ZBTB42*, *ETV1*, *FOS*, *RBPJ*, *NR4A1*, *SIX4*, *ID3*, *ATF3*), as well as the term fat cell differentiation and six associated TF genes (*ATF2*, *NR4A1*, *TCF7L2*, *EGR2*, *PPARG*, *GLIS1*). Module 5, containing *KLF16*, was enriched for terms ncRNA transcription and positive regulation of the Notch signaling pathway (Figure 3D), which is essential in bone development and metabolism (Liu et al., 2016).

In addition, each of the modules was enriched for GO cellular component (CC) terms of various complexes. Module 2 was enriched for the term ESC/E(Z) complex, a multimeric protein complex that can interact with several noncoding RNAs, a vital gene silencing complex regulating transcription, and an effector of response to ovarian steroids (Dubey et al., 2017) (Supplementary Figure 2A). The modules were enriched for different GO molecular function (MF) terms of receptor binding of various hormones, including androgen, estrogen, glucocorticoid, and thyroid hormone receptor binding (Supplementary Figure 2B), providing more evidence of the crosstalk between bone and gonads (Oury, 2012). By RP analysis, Module 2 was associated with activation of HOX genes during differentiation, aligning with their well-established roles in osteogenic differentiation of MSCs and skeletogenesis (Seifert et al., 2015). Module 5 was associated with signaling by TGF-beta family members, and the SMAD2/SMAD3:SMAD4 heterotrimer regulates transcription (Figure 3E). These results revealed the complex and robust nature of TF regulatory networks during osteogenic differentiation.

### Identification of key regulators in the co-expression network of human osteogenic differentiation

We performed multiscale embedded gene co-expression network analysis (MEGENA) to identify gene co-expression structures (i.e., modules) as well as key network regulators (KNRs) (Song and Zhang, 2015; Holmes et al., 2020). MEGENA was conducted on the RNA-seq data from the samples at the MSC, preOB, and OB differentiation stages and identified 168 gene co-expression modules (FDR ≤ 0.05, Figure 4A; Supplementary Table 4). We then identified potential KNRs that were predicted to modulate a large number of DEGs in the network by key regulator analysis (Zhang et al., 2013; Zhang and Zhu, 2013). Many modules enriched for DEGs and their associated KNRs recapitulated known osteogenic factors. For example, Module M204, comprised of 85 genes, was significantly enriched for down-regulated DEGs (MSC to preOB, fold enrichment = 1.80, adjusted p value = 1.58E-2; preOB to OB, fold enrichment = 2.29, adjusted p value = 1.08E-3) (Figure 4A). More than 9% (8 genes) of the DEGs in M204 are related to bone development and metabolism (Supplementary Table 4). Three (*PHLDA2*, *RHOC*, *PFN1*) out of the eight genes were identified as KNRs (Figure 4B). The expression of *PHLDA2* is associated with skeletal growth and childhood bone mass (Lewis et al., 2012). Knockdown of RHOC dramatically inhibits osteogenesis (Zheng et al., 2019). *PFN1*, a member of the profilin family of small actin-binding proteins, is known to regulate bone formation. The loss of PFN1 function causes early onset of Paget’s disease of bone, a disease with impaired osteoclast and osteoblast differentiation (Scotto di Carlo et al., 2020). We also examined the overlap of the genes in M204 with the genes from genome-wide association studies with significant signals for bone area (B-area), bone mineral density (BMD), and hip geometry (Hsu et al., 2019; Morris et al., 2019; Styrkarsdottir et al., 2019), and found five genes, *CYFIP1*, *MMD*, *PELO*, *PRSS23*, and *SNX13*.

**Figure 4.**
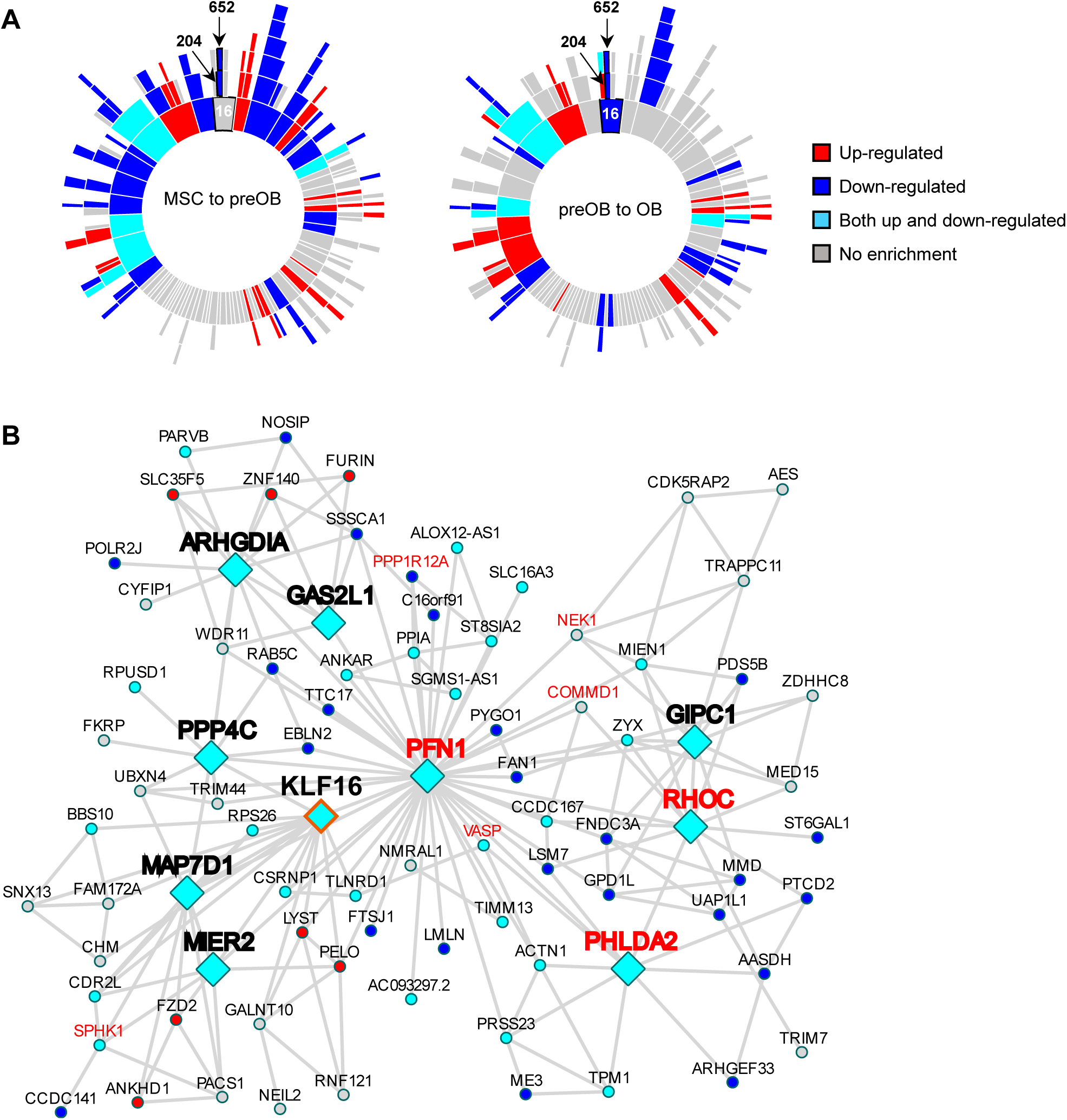
Gene co-expression network in osteogenic differentiation. **(A)** Sunburst plots represent the hierarchy structure of the MEGENA co-expression network constructed on gene expression during the osteogenic differentiation. The structure is shown as concentric rings where the center ring represents the parent modules, and outer rings represent smaller child modules. Subnetwork modules are colored according to the enrichment of differential gene expression between stages (FDR < 0.05; left, MSC to preOB; right, preOB to OB; blue, down-regulated; red, up-regulated; cyan, both up-and down-regulated). The subnetwork branch for Module M204 is outlined and labeled. **(B)** Co-expression network Module M204. Diamonds indicate KNR genes, and circles indicate non-KNR genes. Blue indicates DEGs from MSC to preOB stages, red indicates DEGs from preOB to OB stages, and cyan indicates shared DEGs for both comparisons. Genes known to be related to bone are in red.

As TFs are known to be fundamental to the osteogenic process, we were particularly interested in finding TFs with novel regulatory roles in osteogenesis. We focused on the modules that were significantly enriched for DEGs and contained differentially expressed TFs that were identified as KNRs (Supplementary Table 4). We identified 80 such KNR TFs, of which 60 have unknown biological functions in osteogenesis (Supplementary Table 4). The top 3 up-regulated (*HIF3A*, *ZBTB16*, and *NR2F1*) (Zhu et al., 2014; Onizuka et al., 2016; Manikandan et al., 2018) and the top 2 down-regulated (*TEAD4*, *HMGA1*) (Suo et al., 2020; Wu et al., 2021) KNR TFs have established roles in osteogenesis. The third most down-regulated KNR TF is *KLF16* of Module M204, which previously had little known involvement in osteogenesis (Supplementary Table 4). Thus, based on our cumulative analyses, we hypothesized that *KLF16* plays an important role in osteogenic differentiation.

To further evaluate the expression patterns of the top candidate genes identified through our MEGENA, DEG, and TF-TF network analyses at the cell type level, we employed our previously published single-cell RNA-seq (scRNA-seq) data generated from iPSC-induced cells in osteogenic differentiation (Housman et al., 2022). In that study, gene expression was assessed at two stages, MSCs (Day 0) and osteogenic cells (Day 21) – analogous to our MSC (Day 0) and OB (Day 21) stages for bulk RNA-seq, respectively. Although the differentiation culture conditions differed from this study, we found similar differential expression patterns in a pseudobulk analysis of the scRNA-seq time points for the top 5 gene sets of up- and down-regulated KNR TFs (Supplementary Table 4), TF regulators based on TF-TF regulatory networks (Supplementary Table 3), as well as 5 known osteogenic markers (Supplementary Figure 3; Supplementary Table 5). We further analyzed pseudobulk expression data derived from the five different osteogenic cell types at day 21 and found that most genes of interest showed similar levels of expression among these cell types with the most differences occurring in mature osteocytes, including a notable reduction of RUNX2 expression as expected (Thomas and Jaganathan, 2022) (Supplementary Table 6).

### Regulatory role of *Klf16* in murine osteogenic differentiation

We further focused on *KLF16*, as it is one of the top TFs in the TF regulatory network and a KNR in the co-expression network. *KLF16* is significantly downregulated from the MSC to the OB stage (Figure 5A), suggesting its inhibitory role in osteogenic differentiation. To validate this, we overexpressed *Klf16* in the murine preOB line MC3T3-E1 using lentiviral vector-mediated gene transfer (Figure 5B). We then performed three independent osteogenic differentiation experiments, each with three technical replicates for each differentiation stage (Day 7, 14, and 21). Overexpression of *Klf16* in MC3T3-E1 cells dramatically suppressed osteogenic differentiation at the early stage (Day 7) with reduced ALP activity. At later stages (Day 14 and Day 21), reduced mineralization was detected by alizarin red and von Kossa staining in the *Klf16*-overexpressing cells compared to the control cells (Figure 5C). The inhibitory effect of *Klf16* overexpression on osteogenic differentiation in vitro was observed consistently for each of the experimental and technical replicates.

**Figure 5.**
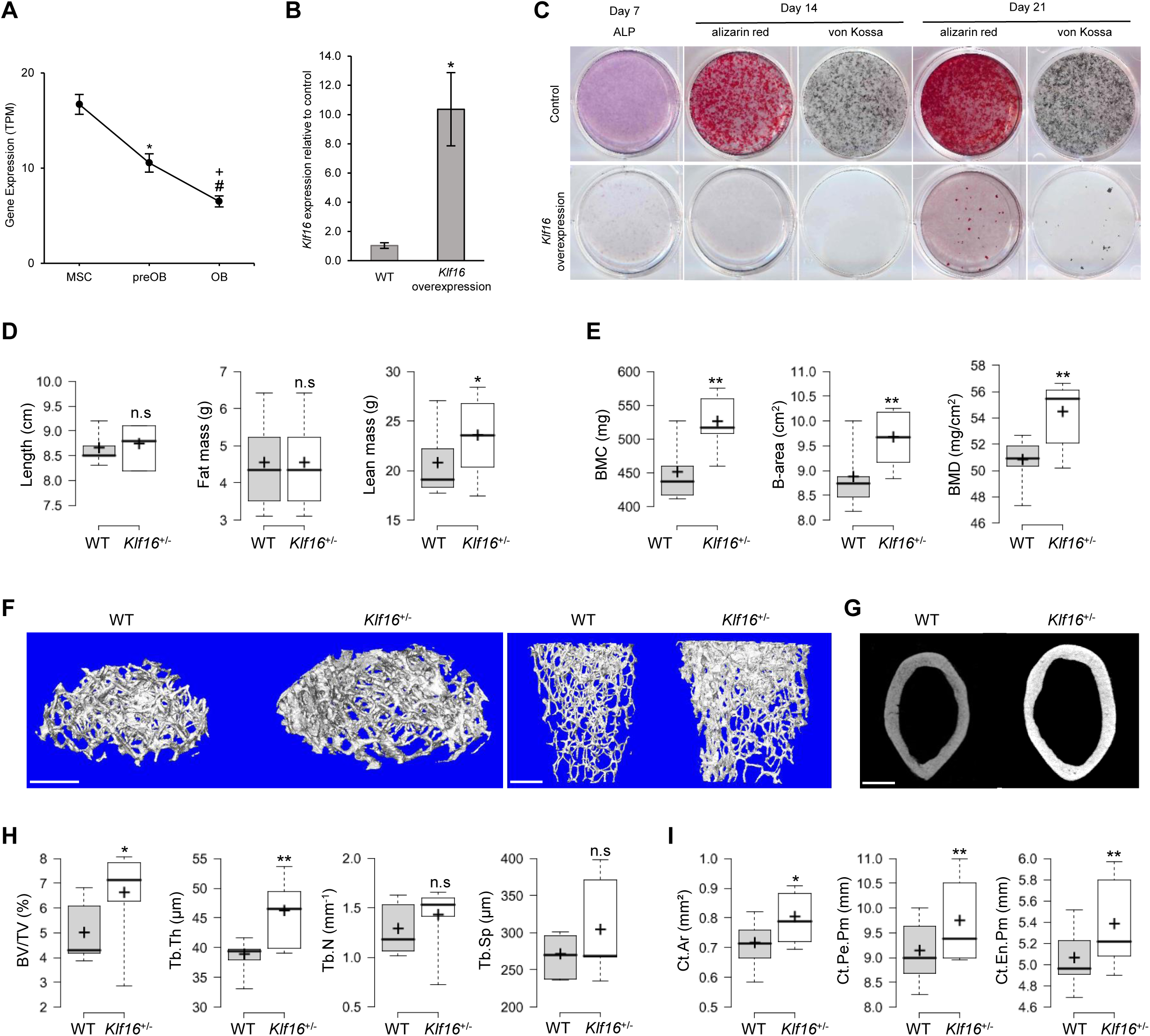
Inhibitory role of *Klf16* in osteogenic differentiation in vitro and in vivo. **(A)** The expression of *KLF16* at three human osteogenic differentiation stages. Data are shown as mean + SEM. * MSC vs. preOB, # preOB vs. OB, or + MSC vs. OB, adjusted p value < 0.001. **(B)** Analysis of *Klf16* expression by RT-qPCR in MC3T3-E1 cells transduced vectors containing either stuffer sequence or *Klf16* cDNA. Data are presented as the mean ± SEM (n=3, unpaired t-test p value < 0.01) (Supplementary Table 7). **(C)** Osteogenic differentiation of MC3T3-E1 cells without or with overexpression of *Klf16*, stained for ALP at Day 7 and alizarin red and von Kossa at Day 14 and Day 21. **(D)** Length, fat mass, and lean mass of wild type (WT) and *Klf16*^+/-^ mice. **(E)** DEXA analysis of whole body (head excluded) bone mineral content (BMC), bone area (B-area), and bone mineral density (BMD) of WT and *Klf16*^+/-^ mice. **(F)** and **(G)** Representative microCT images of distal femur trabecular bone in **(F)** (*left,* top view; *right*, side view) and cortical bone in **(G)** from WT and *Klf16*^+/-^ mice. Scale bar: 1 mm. **(H)** and **(I)** Graphs show trabecular bone volume/tissue volume (BV/TV), trabecular thickness (Tb.Th), trabecular number (Tb.N), and trabecular separation (Tb.Sp) in **(H)**; cortical bone area (Ct.Ar), cortical periosteal perimeter (Ct.Pe.Pm), and cortical endosteal perimeter (Ct.En.Pm) in **(I)**. For **(D)**, **(E)**, **(H)**, and **(I)**, data are presented using BoxPlotR (Spitzer et al., 2014) as the mean ± SEM (WT n = 6, *Klf16*^+/-^ n = 6, 3 males and 3 females for each group, aged 17 weeks, paired t-test, n.s, not significant, * p value < 0.05, ** p value < 0.01).

To explore the role of *Klf16* in bone formation in vivo, we analyzed a *Klf16* knock-out mouse line, generated by deleting a 444 bp segment in exon one using CRISPR technology by the Knockout Mouse Phenotyping Program (KOMP). This mutation deletes the Kozak sequence and the ATG start codon, leaving the final 16 bp of exon one and the splice donor site. We examined adult wild type (WT) and heterozygous (*Klf16^+/-^*) mice, excluding homozygous mice due to preweaning lethality. No significant difference in the overall length between adult *Klf16^+/-^* mice and WT controls was found (Figure 5D). However, adult *Klf16*^+/-^ mice had significantly increased lean mass (total tissue mass – fat mass, including bone mineral content) (p value < 0.05), and no significant change in fat mass relative to WT mice (Figure 5D). Furthermore, there was a dramatic increase in whole body bone mineral content (BMC), bone area, and bone mineral density in *Klf16*^+/-^ mice compared to WT control mice revealed by DEXA scanning (p value < 0.01) (Figure 5E). Microcomputed tomography (microCT) scanning of trabecular bone from adult *Klf16*^+/-^ mice showed significantly increased bone volume fraction (BV/TV) (p value < 0.05) and trabecular thickness (Tb.Th) (p value < 0.01), and no significant difference in trabecular number (Tb.N) or trabecular separation (Tb.Sp) (Figure 5F and 5H). In addition, adult *Klf16*^+/-^ mice showed significantly increased cortical bone area (Ct.Ar) (p value < 0.05), cortical periosteal perimeter (Ct.Pe.Pm) (p value < 0.01), and cortical endosteal perimeter (Ct.En.Pm) (p value < 0.01) compared to WT control mice (Figure 5G and 5I). These findings further support the inhibitory role of KLF16 in bone formation.

Given that bone phenotypes in vivo may be affected by both bone formation and resorption, we analyzed osteoclastogenesis in the *Klf16*^+/-^ mice. The expression of calcitonin receptor (CALCR) as a marker for osteoclasts (Boyce, 2013; Nicholson et al., 1986) was not significantly changed in the epiphysis of the femur, while increased expression was observed in the metaphysis of the *Klf16*^+/-^ mice compared to the controls (Supplementary Figure 4A and 4B). In addition, we examined the expression of receptor activator of NF-kB ligand (RANKL), which binds to receptor activator of NF-kB (RANK) on the surface of osteoclast precursors, promoting their maturation into active osteoclasts and regulating osteoclastogenesis (Khosla, 2001; Boyle et al., 2003). In the femurs of the *Klf16*^+/-^ mice, the expression of RANKL did not show a significant change compared to control mice (Supplementary Figure 4C and 4D). These results suggest that osteoclastogenesis is not decreased in the *Klf16*^+/-^ mice; therefore, increased bone mineral content and density in the mutant mice is more likely attributed to enhanced bone formation rather than reduced resorption by osteoclasts.

## Discussion

Understanding the cellular and molecular mechanisms underlying both normal and pathological bone formation requires insight into the global transcriptional changes during osteogenic differentiation. To investigate these changes, we utilized human iPSC-derived MSCs as our model system. By profiling RNA expression during in vitro osteogenic differentiation, we observed that the transcriptome profiles of our iPSC-derived MSCs and OBs were consistent with those of their primary counterparts, which suggests the potential applicability of our differentiation system to model in vivo osteogenesis. Our differential gene expression analyses provided insights into transcriptional signatures for each osteogenic stage, characterized by changes in expression levels of numerous coding and noncoding genes. Our systems biology approach revealed complex networks of TFs underlying osteogenic differentiation. TF networks in mammalian cells can be built by combining knowledge of TF binding events within TF genes and functional analysis (Wilkinson et al., 2017). Our research unveiled a multifaceted osteogenic differentiation TF network, segmented into five interactive modules. These modules, consistent with previous findings on functional modularity (Dittrich et al., 2008), showcased specific biological functions. For instance, Module 3 underscores TFs associated with fat cell differentiation, suggesting a balance between osteogenesis and adipogenesis within bone structures. This resonates with recent studies emphasizing the role of a network of stem cell TFs in osteogenesis, which act as repressors of adipogenesis (Rauch et al., 2019). Additionally, this module identifies TFs involved in muscle differentiation, indicating overlapping regulatory mechanisms for both bone and muscle development. This concept is further supported by recent discoveries linking mesenchymal progenitors in muscle to bone formation (Julien et al., 2021). This connectivity endorses known interactions between signaling pathways like TGF-β/BMP and Wnt (Hernández-Vega and Camacho-Arroyo, 2021; Thomas and Jaganathan, 2022). Overall, these findings reveal intricate TF regulatory networks and pathway interactions in osteogenic differentiation.

In addition, we explored the gene co-expression network, highlighting potential gene modules and key regulators that play an important role in differentiation. We experimentally validated the regulatory role of *KLF16* from Module M204 as a proof-of-concept that our dataset provides a resource for the discovery of novel mechanisms in osteogenic differentiation. Module M204 is enriched for DEGs, of which many are related to bone formation, including KNR genes *PFN1*, *PHLDA2*, and *RHOC*. We then focused on the KNR *KLF16*, which was not previously associated with bone formation, and investigated its involvement in osteogenic differentiation. Overexpression of *Klf16* in vitro reduced ALP activity and matrix mineralization during MC3T3-E1 differentiation. Conversely, heterozygous *Klf16* knockout mice showed altered bone characteristics, including increases in whole body BMD, femoral bone volume to total tissue volume, trabecular bone thickness, and cortical bone area, suggesting KLF16’s regulatory function in inhibiting osteogenic differentiation.

KLF16, a C2H2-type zinc finger transcription factor, has affinity for GC and GT boxes, and displaces TFs SP1 and SP3 in neural settings (Hwang et al., 2001; Wang et al., 2016). These TFs, when attached to certain promoters, play roles in osteogenic differentiation (Le Mée et al., 2005). Furthermore, KLF16 targets specific metabolic genes, including estrogen-related receptor beta (ESRRB) and PPARA (Daftary et al., 2012; Sun et al., 2021). Our TF regulatory network analysis indicates that *KLF16* is associated with Notch signaling and TGFβ/BMP-SMAD signaling pathways, and regulation of ncRNA transcription. This paints a broad canvas of potential regulatory mechanisms for KLF16 in bone dynamics. Furthermore, other members from the KLF family, especially *KLF4* (Module 1), *KLF5* (Module 5), and *KLF15* (Module 3) (Supplementary Table 4), significantly influence skeletal development (Shinoda et al., 2008; Song et al., 2017; Li et al., 2021; Yu et al., 2021; Zakeri et al., 2022). *KLF5*, co-existing with *KLF16* in Module 5, curtails osteogenesis by inhibiting β-catenin through DNMT3B-induced hypermethylation (Li et al., 2021). KLF4’s modulation of osteogenic differentiation hinges on the BMP4-dependent pathway, resulting in reduced osteoblast numbers in the deficient mice (Yu et al., 2021). Meanwhile, KLF15 enhances chondrogenic differentiation via the TF SOX9 promoter (Song et al., 2017). The roles of KLF6 and KLF9 (Modules 3 and 2) in bone formation remain less defined (Zakeri et al., 2022). Collectively, these findings underline the KLF family’s complex involvement in osteogenesis.

Interestingly, *KLF16* was upregulated in the MSCs of elderly patients suffering from osteoporosis compared to age-matched controls in a previous study (Benisch et al., 2012). A following study identified KLF16 as one of the key TFs whose targets were enriched from the DEGs in the MSCs of the osteoporosis patients versus controls (Liu et al., 2019). The inhibitory role of KLF16 in osteogenic differentiation identified in our study supports the hypothesis that the overexpression of *KLF16* might contribute to osteoporosis, and that KLF16 could therefore be a potential therapeutic target.

Taken together, the differentiation of iPSC-derived MSCs to OBs provides a valuable model for investigating gene expression and regulatory networks in osteogenic differentiation. Our study sheds light on the intricate, layered, and dynamic regulation of the transcriptomic landscape in osteogenic differentiation and offers a foundational resource for further exploration of normal bone formation and the mechanisms driving pathological conditions. Furthermore, this experimental model might facilitate therapeutic research, with potential applications in the treatment of conditions such as osteoporosis.

## Methods

### Human subjects

The twenty human subjects included in this study were in good general health. The study protocols and informed consent were approved by the Institutional Review Board (IRB) of Stanford University and the IRB of the Icahn School of Medicine at Mount Sinai. Each subject gave written informed consent for study participation. Complete subject demographic data can be found in Supplementary Table 1.

### Animal care and use

The *Klf16*^+/-^ mouse strain C57BL/6NJ-Klf16em1(IMPC)J/Mmjax was generated on the C57BL/6NJ genetic background by the Knockout Mouse Phenotyping Program (KOMP) at The Jackson Laboratory (Stock# 032653) using CRISPR technology. Procedures using mice were in compliance with animal welfare guidelines mandated by the Institutional Animal Care and Use Committee (IACUC) and Institutional Biosafety Committee (IBC) at The Jackson Laboratory. Genotyping was performed by real-time PCR of DNA extracted from tail biopsies. A detailed protocol can be found on The Jackson Laboratory website.

### Cell lines

MC3T3-E1 Subclone 4 cell line was purchased from ATCC (CRL-2593, sex undetermined). Irradiated mouse embryonic fibroblasts (MEFs) were purchased from GlobalStem (GSC-6001G). Human iPSC lines were generated in this study. The cells were maintained at 37°C with 5% CO2.

### Peripheral blood mononuclear cell (PBMC) and dermal fibroblast isolation

PBMCs were isolated as we previously described (Carcamo-Orive et al., 2017). Briefly, blood was drawn in tubes containing sodium citrate anticoagulant solution (BD Vacutainer CPT mononuclear cell preparation tube, 362760). PBMCs were separated by gradient centrifugation. The cells were then frozen in RPMI medium (MilliporeSigma, R7388) supplemented with either 12.5% human serum albumin and 10% DMSO (MilliporeSigma, D2438) or 10% fetal bovine serum (FBS) (MilliporeSigma, F4135) and 10% DMSO, and stored in liquid nitrogen for later use. The detailed protocol for dermal fibroblast isolation can be found in our previous publication (Schaniel et al., 2021). Briefly, a skin sample was taken from each of the clinically healthy subjects using a 3 mm sterile disposable biopsy punch (Integra Miltex, 98PUN3-11). Each sample was cut into smaller pieces and placed into gelatin-coated tissue culture dishes with DMEM (Thermo Fisher Scientific, 10567022) supplemented with antibiotics, 20% FBS, non-essential amino acids (Thermo Fisher Scientific, 11140050), 2mM L-glutamine (Thermo Fisher Scientific, 25030081), 2 mM sodium pyruvate (Thermo Fisher Scientific, 11360070), and 100 μM 2-mercaptoethanol (MP Biomedicals, C194705) to establish fibroblast lines. Fibroblasts were harvested using TrypLE Express (Thermo Fisher Scientific, 12605010) and passaged at a 1 to 4 split ratio. Fibroblasts were cryopreserved in 40% DMEM, 50% FBS, and 10% DMSO.

### iPSC generation

Erythroblast protocol: The detailed erythroblast reprogramming process can be found in our previous publication (Carcamo-Orive et al., 2017). Briefly, PBMCs were thawed, and the erythroblast population was expanded for 9–12 days until approximately 90% of cells expressed CD36 and CD71. These expanded erythroblasts were reprogrammed by transduction with Sendai viruses (SeV) expressing factors OCT3/4, SOX2, KLF4, and c-MYC using the CytoTune™-iPS 2.0 Sendai Reprogramming kit (Thermo Fisher Scientific, A16518) according to manufacturer’s protocol.

Fibroblast protocol: The detailed fibroblast reprogramming process can be found in our previous publication (Schaniel et al., 2021). Briefly, mycoplasma-free fibroblasts at passage number 3-5 were reprogrammed using the mRNA reprogramming Kit (Stemgent, 00-0071) in combination with the microRNA booster kit (Stemgent, 00-0073) according to the manufacturer’s protocol.

### iPSCs expansion and characterization

To establish iPSCs, the cells were cultured on Matrigel (BD Bioscience, 354230-10, or Corning, 254248) in feeder-free conditions and maintained in mTeSR1 (Stem Cell Technologies, 05850). The iPSCs were passaged every 5-7 days after clones were picked. To ensure the high quality of the iPSC lines used in this study, those lines generated with Sendai virus first underwent virus clearance by continuous passaging. To measure the loss of SeV in the iPSCs generated from erythroblasts with SeV, quantitative RT-PCR was performed according to the SeV reprogramming kit manufacture protocol. For quality control, G-banded karyotyping, ALP staining, iPSC pluripotency marker immunocytochemistry, and embryoid body formation analysis (Yaffe et al., 2016) were performed. Differentiating colonies were routinely eliminated from the cultures. All the lines included in the current study had normal karyotypes, exhibited human embryonic stem cell morphology, and expression of ALP and pluripotency markers NANOG, SOX2, OCT4, SSEA4, and TRA-1-60. Further, embryoid body (EB) formation and differentiation assays demonstrated their competence for differentiation into all three germ layers. We excluded samples where any of these characterizations were abnormal (Daley et al., 2009; Yaffe et al., 2016).

### Mycoplasma quality control

Mycoplasma testing of the cells was performed intermittently during culture. Cultures were grown in the absence of antibiotics for at least three days before testing. The culture medium or the DNA harvested from the cultures were tested for mycoplasma contamination using a MycoAlert Mycoplasma Detection Kit (Lonza, LT07-418) or e-Myco PLUS PCR Detection Kit (BOCA scientific, 25237).

### ALP staining

Cells cultured on plates were briefly washed with phosphate-buffered saline (PBS), fixed in 4% PFA for 3 minutes at room temperature, and then stained with Alkaline phosphatase kit II (Stemgent, 00-0055) according to the manufacturer’s instructions.

### iPSC embryoid body analysis

iPSC embryoid body formation and characterization were performed as previously described with modifications (Lin and Chen, 2008). Briefly, iPSCs were cultured in mTeSR1 medium on Matrigel-coated plates, and the medium was changed daily. On the day of EB formation, when the cells grew to 60-80% confluence, cells were washed once with PBS and then incubated in Accutase for 8-10 minutes to dissociate colonies to single cells and resuspended with mTeSR1 containing 2 µM Thiazovivin. To form self-aggregated EBs, single iPSCs were transferred into an ultra-low attachment plate (Corning, CLS3471) via 1:1 or 1:2 passage. EBs were aggregated from iPSCs for three days, then were transferred to 0.1% gelatin-coated 12 well plates and maintained in embryoid body medium (DMEM/F12 supplemented with 10% FBS, 2 mM L-glutamine, 0.1 mM non-essential amino acids, and 0.1 mM 2–mercaptoethanol) for another ten days for the spontaneous generation of three germ layers with medium changed every other day. Fluorescent immunostaining was performed to detect the expression of markers of the three germ layers.

### Fluorescence immunocytochemistry

The cells cultured on plates were briefly washed with PBS, fixed in 4% paraformaldehyde (PFA) in PBS for 15 minutes, washed with PBS three times, and then permeabilized with 0.2% Triton in PBS for 15 mins. The cells were then washed with PBS three times prior to blocking with 1% bovine serum albumin (BSA) (MilliporeSigma, A8412) in PBS for 1 hour at room temperature and incubating with primary antibody 1:100 to 1:900 dilutions in 0.1% BSA in PBS for overnight at 4°C. Washing three times was conducted with PBS followed by incubation with a corresponding secondary antibody in 1:500 dilution in 0.1% BSA for 1 hour at room temperature. After washing with PBS, 1:50,000 diluted Hoechst (Thermo Fisher Scientific, H1398) was added to stain nuclei for 5 minutes, followed by washing with PBS three times and imaging. The following primary antibodies were applied to detect the expression of iPSC markers: anti-TRA-1-60 (Invitrogen, 41-1000, 1/100), anti-SSEA4 (Invitrogen, 41-4000, 1/200), anti-NANOG (Abcam, ab109250, 1/100), anti-OCT-4 (Cell Signaling Technology, 2840S, 1/400), and anti-SOX2 (Abcam, ab97959, 1/900). To detect the expression of the markers of three germ layers, anti-AFP (Agilent, A0008, 1/100) for endoderm, anti-α-SMA (MilliporeSigma, A5228, 1/200) for mesoderm, and anti-TUBB3 (MilliporeSigma, T2200, 1/200) for ectoderm were used. All the Alexa-conjugated secondary antibodies used were from Invitrogen: anti-mouse Alexa Fluor 488 (A-11029), anti-rabbit Alexa 488 (A-21206), and anti-rabbit Alexa 594 (A-11037).

### G-banding karyotyping

Karyotype analysis was performed on all iPSC lines by the Human Genetics Core facility at SEMA4. iPSCs were cultured on Matrigel-coated T25 flasks before karyotyping and reached approximately 50% confluence on the day of culture harvest for karyotyping. Twenty cells in metaphase were randomly chosen, and karyotypes were analyzed using the CytoVision software program (Version 3.92 Build 7, Applied Imaging).

### Differentiation of iPSCs to MSCs

In vitro differentiation of iPSCs to MSCs was carried out with commercial cell culture media according to the manufacture protocols with modifications. Briefly, iPSC colonies were harvested with Accutase (Innovative Cell Technologies, AT-104) and seeded as single cells on a Matrigel-coated plate in mTeSR1 and supplemented with 2uM Thiazovivin. The next day, TeSR1 medium was replaced with STEMdiff mesoderm induction medium (MIM) (Stem Cell Technologies, 05221) when cells were at approximately 20-50% confluence. Cells were then fed daily and cultured in STEMdiff MIM for four days. On Day 5, the culture medium was switched to MesenCult-ACF medium (Stem Cell Technologies, 05440/8) for the rest of the MSC induction duration. Cells were passaged as necessary using MesenCult-ACF Dissociation Kit (Stem Cell Technologies, 05426). Cells were subcultured onto Matrigel at the first passage to avoid loss of the differentiated cells before tolerating the switch to MesenCult-ACF attachment substrate (Stem Cell Technologies, 05448). For passage two or above, cells were subcultured on MesenCult-ACF attachment substrate. After three weeks of differentiation, the cells were sorted for the CD105 (BD Biosciences, 561443, 1/20) positive and CD45 (BD Biosciences, 555483, 1/5) negative MSC population (Giuliani et al., 2011; Kang et al., 2015; Sotiropoulou et al., 2006) using BD FACSAria II in the Mount Sinai Flow Cytometry Core Facility, and then expanded in MSC medium consisting of low-glucose DMEM (Gibco, 10567022) containing 10% FBS (Sotiropoulou et al., 2006). After expansion, a few MSC lines were further examined by BD CantoII for expression of other MSC positive surface markers CD29 (Thermo Fisher Scientific, 17– 0299, 1/20), CD73 (BD Biosciences, 561254, 1/20), CD90 (BD Biosciences, 555595,1/ 20), absence of other negative surface markers CD31 (BD Biosciences, 561653, 1/20), CD34 (BD Biosciences, 560940, 1/5), and retention of marker CD105 positivity and CD45 negativity as well.

### Fluorescence activated cell sorting (FACS) and analysis

Single cells were washed with FACS buffer (PBS supplemented with 1% FBS and 25mM HEPES (Thermo Fisher Scientific, 1688449)) twice, resuspended to a concentration of 1X10^7^ cells/ml in ice-cold FACS buffer, and incubated with the above conjugated antibodies for 30 minutes at 4° in the dark. Stained cells were then washed with FACS buffer three times before FACS or flow cytometry analysis. Flow cytometry data were analyzed with BD FACSDiva software.

### Differentiation of iPSC-derived MSCs to osteoblasts

iPSC-derived MSCs were plated in a 6-well plate at a density of 3X10^3^ cells/cm^2^ in MSC medium. After three days, the culture medium was switched to osteogenic differentiation medium (α MEM (Thermo Fisher Scientific, A1049001) supplemented with 10% FBS, 1% non-essential amino acids, 0.1 µM dexamethasone (MilliporeSigma, D4902), 10 mM β-glycerophosphate (MilliporeSigma, G9422), and 200 µM ascorbic acid (MilliporeSigma, A4544)) and maintained in this medium for 21 days (Barberi et al., 2005; Lee et al., 2015; Pittenger et al., 1999). The culture medium was changed every 2-3 days. Cells were harvested for total RNA isolation and RNA sequencing at Day 0, Day 7, and Day 21 of differentiation, as indicated in the main text. ALP staining, and alizarin red staining and von Kossa staining, were employed to examine bone ALP expression and mineralization, respectively.

### Alizarin red staining and von Kossa staining

Mineralization of osteoblast extracellular matrix was assessed by both alizarin red staining and von Kossa staining. Cells were washed briefly with PBS, fixed with 4% PFA for 15 minutes, and washed with deionized distilled water three times. For alizarin red staining, fixed cells were incubated in alizarin red stain solution (MilliporeSigma, TMS-008-C) with gentle shaking for 30 minutes, followed by washing with water four times for 5 minutes each with gentle shaking to remove non-specific alizarin red staining. For von Kossa staining, fixed cells were incubated in 5% silver nitrate solution (American Master Tech Scientific, NC9239431) while exposed to UV light for 20 minutes. Unreacted silver was removed by incubating in 5% sodium thiosulfate (American Master Tech Scientific, NC9239431) for 5 minutes, followed by washing with water twice. Mineralized nodules were identified as red spots by alizarin red staining and dark brown to black spots by von Kossa staining.

### RNA-seq library preparation and sequencing

For total RNA isolation, cells were washed with PBS and harvested for RNA extraction using miRNeasy mini kit (QIAGEN, 217004) according to the manufacturer’s instruction. Illumina library preparation and sequencing were conducted by the Genetic Resources Core Facility, Johns Hopkins Institute of Genetic Medicine (Baltimore, MD). RNA concentration and quality were determined using NanoDrop Spectrophotometer (Thermo Scientific, DE). RNA integrity (RNA Integrity Number, RIN) was verified using Agilent BioAnalyzer 2100 and the RNA Nano Kit prior to library creation. Illumina’s TruSeq Stranded Sample Prep kit was used to generate libraries. Specifically, after ribosomal RNA (rRNA) depletion, RNA was converted to cDNA and size selected to 150-200 bp in length, then end-repaired and ligated with appropriate adaptors. Ligated fragments were subsequently size-selected and underwent PCR amplification techniques to prepare the libraries with a median size of 150 bp. Libraries were uniquely barcoded and pooled for sequencing. The BioAnalyzer was used for quality control of the libraries to ensure adequate concentration and appropriate fragment size. Sequencing was performed on an Illumina HiSeq 2500 instrument using standard protocols for paired-end 100 bp sequencing. All the samples were processed in one batch.

### RNA-seq pre-processing and gene differential expression analyses

Illumina HiSeq reads were processed through Illumina’s Real-Time Analysis (RTA) software generating base calls and corresponding base call quality scores. CIDRSeqSuite 7.1.0 was used to convert compressed bcl files into compressed fastq files. After adaptor removal with cutadapt (Martin, 2011) and base quality trimming to remove 3′ read sequences if more than 20 bases with Q ≥ 20 were present, paired-end reads were mapped to the human GENCODE V29 reference genome using STAR (Dobin et al., 2013) and gene count summaries were generated using featureCounts (Liao et al., 2014). Raw fragment (i.e., paired-end read) counts were then combined into a numeric matrix, with genes in rows and experiments in columns, and used as input for differential gene expression analysis with the Bioconductor Limma package (Ritchie et al., 2015) after multiple filtering steps to remove low-expressed genes. First, gene counts were converted to FPKM (fragments per kb per million reads) using the RSEM package (Li and Dewey, 2011) with default settings in strand-specific mode, and only genes with expression levels above 0.1 FPKM in at least 20% of samples were retained for further analysis. Additional filtering removed genes less than 200 nucleotides in length. Finally, normalization factors were computed on the filtered data matrix using the weighted trimmed mean of M values (TMM) method, followed by voom (Law et al., 2014) mean-variance transformation in preparation for Limma linear modeling. The limma generalized linear model contained fixed effects for sex (male/female), and cell source (fibroblast/erythroblast) and a random effect term was included for each unique subject. Data were fitted to a design matrix containing all sample groups, and pairwise comparisons were performed between sample groups (i.e., MSC stage, preOB stage, and OB stage). eBayes adjusted p values were corrected for multiple testing using the Benjamin-Hochberg (BH) method (Benajmin and Hochberg, 1995) and used to select genes with significant expression differences (adjusted p value < 0.05).

### Integration of RNA-Seq data of human primary MSCs and OBs, iPSCs, and tissues in GTEx

GTEx gene expression data (version 7) and metadata were downloaded from the GTEx website (Ardlie et al., 2015). Raw counts were extracted for the brain, heart, kidney, liver, lung, muscle, nerve, ovary, testis, and thyroid tissues. After filtering the lowly expressed genes (1 CPM in less than 20% samples), normalization factors were computed using the weighted trimmed mean of M values (TMM) method, followed by the voom mean-variance transformation (Law et al., 2014). The Human iPSC dataset was from our previous publication, deposited in the GEO database with accession number GSE79636 (Carcamo-Orive et al., 2017). Two datasets for each primary human MSC and OB cell type were collected from the GEO accessions GSE94736 (Roforth et al., 2015) and GSE118808 (Kaczynski et al., 2003; Ma et al., 2019), and GSE55282 (Rojas-Peña et al., 2014) and GSE75524 (Al-Rekabi et al., 2016), respectively. All the datasets were filtered and normalized in the same way as the GTEx data. Genes from all the datasets were intersected to obtain shared genes. Gene expression levels of the shared genes were extracted from all the datasets. Principal component analysis (PCA) was performed on the extracted and combined gene expression matrix. Multi-dimensional scaling was used to visualize the top two principal components.

### Gene ontology and Reactome pathway enrichment analyses

Gene ontology (GO) biological process (BP), molecular function (MF), and cellular component (CC), and Reactome pathway enrichment analyses were performed using the PANTHER classification system (www.pantherdb.org) (Mi et al., 2019a). The statistical overrepresentation test in PANTHER was fulfilled by Fisher’s exact test together with FDR multiple test correction to identify enriched GO categories and Reactome Pathways among the input genes relative to the indicated reference list as stated in figure legends (The Gene Ontology Consortium, 2019; Mi et al., 2019b). Enrichment tests were filtered using FDR < 0.05.

### Transcription factor network analysis

The ChIP-seq and ChIP-exo datasets of DNA binding peaks of human transcription regulators (TRs), where the overwhelming majority are TFs, were downloaded from ReMap 2020 (Chèneby et al., 2020). The genomic coordinates of the transcription start sites of human genes were extracted from Ensembl Human Genes GRCh37.p13, and then applied BedTools v2.3.0 to identify the target TR genes of a TR by checking the intersection of the latter TR’s binding peaks and the 2kb window before and after the transcription start sites of the target TR genes (Quinlan and Hall, 2010). The Python package scipy.stats.pearsonr was used to calculate the correlation coefficient between the expression of TR genes (Hao et al., 2015; Jiang et al., 2014; Virtanen et al., 2020; Zhu et al., 2015). The regulatory relationship between TRs was predicted using two criteria: (1) the binding site peaks of a TR are within the distance of 2 kb upstream or downstream of the known transcription start sites of the target TRs; and (2) the absolute value of the correlation coefficient between the TR genes is greater than 0.6 and adjusted p value is less than 0.05 (Camacho et al., 2005). Gephi 0.9.2 was used to generate and visualize the regulatory network, where each node was defined as a TR gene, and two nodes were connected by an edge when ReMap data demonstrated regulation between the two TRs, and our RNA-seq data also showed expression correlation between them. Furthermore, the network community detection hierarchical algorithm was applied to determine relationships between subsets of the whole network and define modules. Betweenness centrality for each of the nodes was calculated and then used to rank node sizes (Bastian et al., 2009).

### Gene co-expression network analysis and identification of key network regulators

Gene co-expression network was constructed using the R package MEGENA v1.3.7 (Song and Zhang, 2015). The same filtered, normalized, and covariate-adjusted gene expression data matrix used in the differential gene expression analysis was used as input for MEGENA. Specifically, the Pearson correlation was used to calculate significant correlations between gene pairs among the 60 samples. With a cutoff of 0.05 FDR, significant correlations were identified by 100 permutations of the gene expression matrix. Next, planar-filtered network construction and multi-scale clustering analysis were performed. Finally, significant modules were identified at a 5% FDR with 100 times network permutations. Modules of smaller than 50 genes or larger than 5,000 genes were excluded from the downstream analyses. Enrichment analysis was performed between the modules and DEG signatures between MSC and preOB stages and between preOB and OB stages through Fisher’s exact test. BH procedure was applied to the p values to correct for multiple-testing problem (Benjamini and Hochberg, 1995). Modules were visualized using the Cytoscape (Otasek et al., 2019). Sunburst plots of modules were visualized using the R package *sunburstR* v2.1.5. We used all the MEGENA nodes and edges as input to identify KNRs that were predicted to modulate a large number of downstream DEGs in the network (Zhang et al., 2013; Zhang and Zhu, 2013). For each gene in the MEGENA network, we tested whether the network neighborhood of the gene was enriched with the DEG signature. Specifically, we tested if the nodes within a path length of 6 steps of the candidate KNR gene were enriched for the DEGs using Fisher’s Exact test. The p values were then corrected by the Bonferroni procedure to adjust for multiple comparisons.

### Processing of comparative single-cell pseudobulk data

scRNA-seq data (GEO accession: GSE181744) from our previous publication (Housman et al., 2022), which includes mixtures of iPSC-derived, MSC osteogenic differentiations from different species, was used as a comparative dataset. To maximize the comparative utility of these data, we re-processed the raw scRNA-seq data to obtain pseudobulk expression values for all genes annotated in the human genome and for cell types of interest from the six humans and one human technical replicate included in this dataset. Briefly, reads were processed using standard 10X Genomics Cell Ranger 3.1.0 pipelines (Zheng et al., 2017) that extracted 10X cell barcodes and UMIs and aligned the remaining reads to the human genome (hg38). Human cells and specific cell types of interest (MSCs, osteogenic cells, preosteoblasts, osteoblasts, embedding osteoblasts, mineralizing osteoblasts, and maturing osteocytes) were isolated from the newly processed data using species assignments and cell classifications (Housman et al., 2022). Lastly, single-cell gene count data were consolidated to produce pseudobulk expression values for each unique grouping of individual, replicate, and cell classification. Specifically, these pseudobulk data were defined as the sum of raw single-cell UMI counts within each individual-replicate for a given cell classification. Computational scripts for these processing steps can be found on GitHub at https://github.com/ghousman/human-skeletal-scRNA. Pseudobulk data were filtered and normalized as described above.

### Generation of *Klf16*-overexpressed MC3T3-E1 cell line

Lentiviral transduction of MC3T3-E1 Subclone 4 cells (ATCC, CRL-2593) was performed as we previously described with modifications (Holmes et al., 2020). Cells were infected by incubating in lentivirus-containing cell culture medium at a multiplicity of infection (MOI) of 100 in the presence of 6 μg/ml polybrene for 24 hours. The selection was performed with 2 µg/ml puromycin for 12 days until EGFP expression was observed in 100% of cells, no further cell death was observed, and no live cells remained in the non-transduction negative control dish. Selected cells were expanded and cryopreserved. EGFP expression was monitored routinely, and additional selection was performed when necessary.

### MC3T3-E1 osteoblast differentiation assay

Osteoblast differentiation of MC3T3-E1 transfected with lentiviral vectors containing either *Klf16* cDNA or a 300bp nonfunctional stuffer sequence was performed as follows: cells were plated in 6-well plates at a density of 3X10^3^ cells/cm^2^ in MC3T3-E1 maintenance medium (αMEM supplemented with 10% FBS). After three days, the culture medium was switched to osteogenic differentiation medium (Holmes et al., 2020), and cells were maintained in this medium, which was replenished every two days for 21 days. Triplicate experiments, each having triplicate wells, were carried out for both control and *Klf16*-overexpressing cells.

### RT-qPCR

The RT-qPCR method was used to assess gene expression in stable lentiviral MC3T3-E1 cell lines. When cells were plated in 6 well plates for osteogenic differentiation as described above, one parallel well was used for RNA isolation after being cultured in MC3T3-E1 maintenance medium for three days before switching to osteogenic differentiation medium. Total RNA was extracted with the RNeasy Kit (Qiagen, 74106) according to the manufacturer’s protocol. cDNA was synthesized with AffinityScript One-Step RT-PCR Kit (Agilent, 600188). Each cDNA sample was amplified in triplicate using the SYBR Green and Platinum Taq polymerase (Thermo Fisher Scientific, S7567) on a 7900HT Real-Time PCR instrument (Thermo Fisher Scientific, 10966). The housekeeping gene *Actb* was used as the reference. mRNA relative expression was calculated by the ΔΔCt method. qPCR primers used are listed in Supplementary Table 7.

### Whole body bone mineral density

Bone mineral density (BMD) was scanned with a Dual Energy X-ray Absorptiometry (DEXA) Analyzer (Lunar, Piximus II, GE Medical System). Mouse weight and length were measured (nose to the beginning of tail) before scanning. Each unconscious mouse was placed in the DEXA analyzer. A scout-scan was performed. The analysis was conducted on a whole body scan, excluding the head. The mouse was removed once the image was captured and placed on a heated mat set at 37°C in a cage and closely monitored until consciousness was regained.

### Microcomputed tomography

Femurs were isolated by removing attached soft tissues, fixed in 4% PFA at 4°C, and imaged with microCT scan (Skyscan 1172a, Skyscan, Belgium) at 50 kV and 200 µA using a 0.5 mm aluminum filter and an image voxel size of 4.5 µm isotropic. Images were captured every 0.7°, with 8× averaging, through 180° rotation of each bone and reconstructed using Skyscan NRecon software. Imaging analysis of the metaphyseal regions of each femur was performed by first determining a reference point of the most proximal slice where the growth plate had begun to disappear. Offsets of 101 slices and 501 slices from the point of reference in the growth plate were used for trabecular and cortical analyses, respectively, with Skyscan CTAn software. Volume visualization was performed in Avizo 2020.2 (Thermo Fisher Scientific). Data were reported in the standard format used by the American Society for Bone and Mineral Research (ASBMR) (Parfitt et al., 1987). Each volume was then segmented to isolate osteological material using a regularized deep network (RDN)-based image segmentation algorithm (Yazdani et al., 2020). The segmented images were thresholded and masked in Medtool 4.4 (DPI e.U, Medtool) to isolate trabecular bone from cortical bone (Gross et al., 2014). The external mask (cortical bone) was subtracted from the volume, leaving only the internal mask (trabecular bone) and air. Bone volume fraction (BV/TV), trabecular thickness (Tb.Th), trabecular number (Tb.N), and trabecular separation (Tb.Sp) were calculated from the trabecular bone volumes in 3D using 7 mm sampling spheres on a background grid with 3.5 mm spacing following DeMars et al. (2021) in Medtool. Cortical bone variables cortical area (Ct.Ar), cortical periosteal perimeter (Ct.Pe.Pm), and cortical endosteal perimeter (Ct.En.Pm) were then calculated in the BoneJ extension of ImageJ (Doube et al., 2010) for each volume.

### Immunohistochemistry

Femurs from two male *Klf16^+/-^* mice and two male WT mice at the age of 18 weeks were fixed and prepared for paraffin section. Sections were deparaffinized in xylene and rehydrated through a gradient of ethanol (100%, 95%, and 70%, 5 minutes each) to water. Slides were incubated in 0.2% Triton X-100 for 5 min. After washing in PBS, endogenous fluorescence was blocked using TrueBlack Plus Lipofuscin Autofluorescence Quencher (1:40, Biotium, 23014) for 20 min, and then blocked with an Animal-Free Blocker (Vector Laboratories, SP-5035-100) for 1 hour at RT. The primary antibody anti-CALCR (1:50, Bioss, BS-0124R), or anti-RANKL (1:50, Abcam, ab216484), was applied and incubated overnight at 4°C. After washing in PBS, secondary antibody (Alexa Fluor 488 chicken anti-Rabbit IgG, 1:500, Invitrogen, A-21441) was applied and incubated for 1 hour at room temperature. Slides were then washed in PBS, stained with DAPI, and mounted with Antifade Mounting Media (Vector Labs, H-1700-10). Images were acquired with a Nikon T1-SM microscope.

### Statistical analyses

Statistical analyses were performed with Microsoft Excel using Student’s t-test to create graphs displaying mean values with error bars corresponding to the standard error of the mean (SEM) for *Klf16* in vitro overexpression and animal experiments. p value < 0.05 was considered significant. Sample sizes are indicated in figure legends. Image analyses were performed with ImageJ (National Institutes of Health). IHC data were quantified and visualized with GraphPad Prism 10 (GraphPad Software).

### Data and code availability

The RNA-seq data supporting the findings of this study are deposited in Gene Expression Omnibus (GEO) with accession number GSE200492.

## Acknowledgements

We thank Paige Cundiff for support in iPSC derivation, Arvind Babu for chromosome analyses, Xuqiang Qiao for assistance in the Dean’s Flow Cytometry Core, and Jacqui White and Arie Mobley of The Jackson Laboratory Center for Biometric Analysis for the microCT imaging and DEXA scanning of the *Klf16* mouse line. This work was supported in part through the computational resources and staff expertise provided by Scientific Computing at the Icahn School of Medicine at Mount Sinai. The research reported in this paper was supported by the Office of Research Infrastructure of the National Institutes of Health under award numbers S10OD026880 and S10OD030463. This work was funded by NIH grants P01HD078233 (Ethylin Wang Jabs) and R01DE029832 (Ethylin Wang Jabs, Susan M. Motch Perrine) and F32AR075397 (Genevieve Housman). The content is solely the responsibility of the authors and does not necessarily represent the official views of the National Institutes of Health.

## Author details

### Ying Ru

Department of Genetics and Genomic Sciences, Icahn School of Medicine at Mount Sinai, New York, NY, 10029, USA

**Contribution:** Conceptualization, Methodology, Validation, Formal analysis, Investigation, Data curation, Writing – original draft, Writing – review & editing, Visualization, Supervision

**Competing interests:** No competing interests declared

### Meng Ma

Mount Sinai Genomics, Sema4, Stamford, CT, 06902, USA

**Contribution:** Conceptualization, Methodology, Validation, Formal analysis, Investigation, Data curation, Writing – original draft, Writing – review & editing, Visualization

**Competing interests:** No competing interests declared

### Xianxiao Zhou

Department of Genetics and Genomic Sciences, Icahn School of Medicine at Mount Sinai, New York, NY, 10029, USA

Mount Sinai Center for Transformative Disease Modeling, Icahn School of Medicine at Mount Sinai, New York, NY, 10029, USA

Icahn Genomics Institute, Icahn School of Medicine at Mount Sinai, New York, NY, 10029, USA

**Contribution:** Methodology, Validation, Formal analysis, Investigation, Data curation, Writing – original draft, Writing – review & editing, Visualization

**Competing interests:** No competing interests declared

### Divya Kriti

Department of Genetics and Genomic Sciences, Icahn School of Medicine at Mount Sinai, New York, NY, 10029, USA

Present address: Department of Biochemistry and Molecular Biology, Faculty of Medicine, The University of British Columbia, Vancouver, BC V6T 2G3, Canada

**Contribution:** Formal analysis, Investigation, Data curation, Writing – review & editing

**Competing interests:** No competing interests declared

### Ninette Cohen

Department of Genetics and Genomic Sciences, Icahn School of Medicine at Mount Sinai, New York, NY, 10029, USA

Present address: Division of Cytogenetics and Molecular Pathology, Zucker School of Medicine at Hofstra/Northwell, Northwell Health Laboratories, Lake Success, NY, 11030, USA

**Contribution:** Investigation, Writing – review & editing

**Competing interests:** No competing interests declared

### Sunita D’Souza

Department of Cell, Developmental and Regenerative Biology, Icahn School of Medicine at Mount Sinai, New York, NY, 10029, USA

Present address: St Jude Children’s Research Hospital, Memphis, TN, 38105, USA

**Contribution:** Methodology - iPSC derivation, Writing – review & editing

**Competing interests:** No competing interests declared

### Christoph Schaniel

Department of Cell, Developmental and Regenerative Biology, Icahn School of Medicine at Mount Sinai, New York, NY, 10029, USA

Department of Medicine, Division of Hematology and Medical Oncology, Tisch Cancer Institute, Icahn School of Medicine at Mount Sinai, New York, NY, 10029, USA

Black Family Stem Cell Institute, Icahn School of Medicine at Mount Sinai, New York, NY, 10029, USA

Mount Sinai Institute for Systems Biomedicine, Icahn School of Medicine at Mount Sinai, New York, NY, 10029, USA

**Contribution:** Methodology - iPSC derivation, Writing – review & editing

**Competing interests:** No competing interests declared

### Susan M. Motch Perrine

Department of Anthropology, Pennsylvania State University, University Park, PA, 16802, USA

**Contribution:** Writing – review & editing, Visualization

**Competing interests:** No competing interests declared

### Sharon Kuo

Department of Biomedical Sciences, University of Minnesota, Duluth, MN, 55812, USA Technological Primates Research Group, Max Planck Institute for Evolutionary Anthropology, Leipzig 04103, Germany.

**Contribution:** Writing – review & editing, Visualization

**Competing interests:** No competing interests declared

### Oksana Pichurin

Department of Clinical Genomics, Mayo Clinic, Rochester, MN, 55905, USA

**Contribution:** Investigation

**Competing interests:** No competing interests declared

### Dalila Pinto

Department of Genetics and Genomic Sciences, Icahn School of Medicine at Mount Sinai, New York, NY, 10029, USA

**Contribution:** Formal analysis, Writing – review & editing

**Competing interests:** No competing interests declared

### Genevieve Housman

Section of Genetic Medicine, Department of Medicine, University of Chicago, Chicago, IL, 60637, USA

Department of Primate Behavior and Evolution, Max Planck Institute for Evolutionary Anthropology, Leipzig, 04103, Germany

**Contribution:** Writing – review & editing

**Competing interests:** No competing interests declared

### Greg Holmes

Department of Genetics and Genomic Sciences, Icahn School of Medicine at Mount Sinai, New York, NY, 10029, USA

**Contribution:** Conceptualization, Writing – review & editing

**Competing interests:** No competing interests declared

### Eric Schadt

Department of Genetics and Genomic Sciences, Icahn School of Medicine at Mount Sinai, New York, NY, 10029, USA

Icahn Genomics Institute, Icahn School of Medicine at Mount Sinai, New York, NY, 10029, USA

**Contribution:** Conceptualization, Methodology, Writing – review & editing

**Competing interests:** No competing interests declared

### Harm van Bakel

Department of Genetics and Genomic Sciences, Icahn School of Medicine at Mount Sinai, New York, NY, 10029, USA

Icahn Genomics Institute, Icahn School of Medicine at Mount Sinai, New York, NY, 10029, USA

Department of Microbiology, Icahn School of Medicine at Mount Sinai, New York, NY, 10029, USA

**Contribution:** Conceptualization, Methodology, Validation, Formal analysis, Investigation, Data curation, Writing – original draft, Writing – review & editing, Visualization, Supervision

**Competing interests:** No competing interests declared

### Bin Zhang

Department of Genetics and Genomic Sciences, Icahn School of Medicine at Mount Sinai, New York, NY, 10029, USA

Mount Sinai Center for Transformative Disease Modeling, Icahn School of Medicine at Mount Sinai, New York, NY, 10029, USA

Icahn Genomics Institute, Icahn School of Medicine at Mount Sinai, New York, NY, 10029, USA

**Contribution:** Conceptualization, Methodology, Validation, Writing – review & editing, Supervision

**Competing interests:** No competing interests declared

### Ethylin Wang Jabs

Department of Genetics and Genomic Sciences, Icahn School of Medicine at Mount Sinai, New York, NY, 10029, USA

Department of Cell, Developmental and Regenerative Biology, Icahn School of Medicine at Mount Sinai, New York, NY, 10029, USA

Department of Clinical Genomics, Mayo Clinic, Rochester, MN, 55905, USA

Department of Biochemistry and Molecular Biology, Mayo Clinic, Rochester, MN, 55905, USA

**Contribution:** Conceptualization, Validation, Writing – review & editing, Supervision, Project administration, Funding acquisition

**Competing interests:** No competing interests declared

### Meng Wu

Department of Genetics and Genomic Sciences, Icahn School of Medicine at Mount Sinai, New York, NY, 10029, USA

Department of Clinical Genomics, Mayo Clinic, Rochester, MN, 55905, USA

Department of Biochemistry and Molecular Biology, Mayo Clinic, Rochester, MN, 55905, USA

**Contribution:** Investigation, Writing – review & editing, Supervision

**Competing interests:** No competing interests declared

---

**Supplementary Figure 1.**
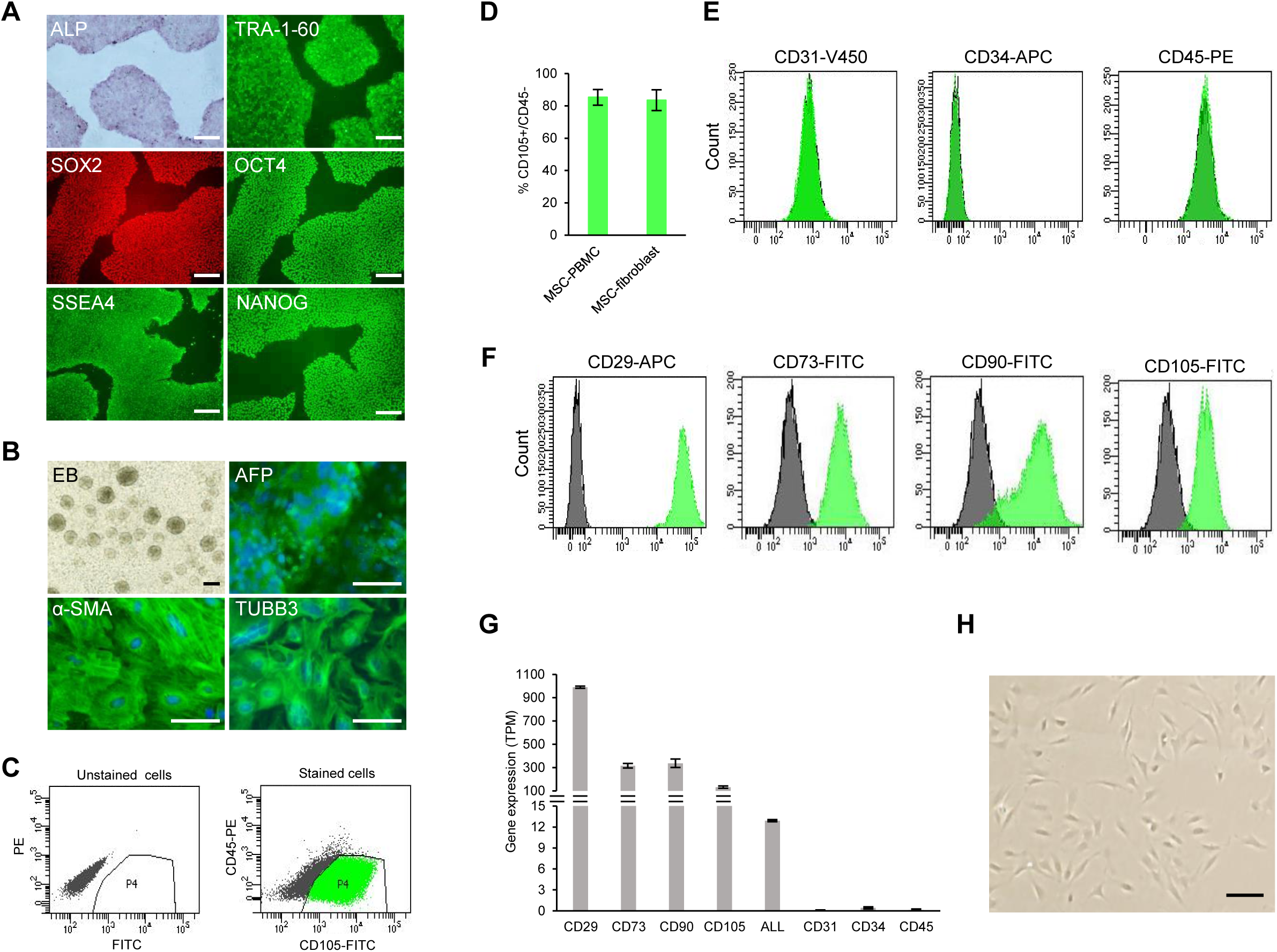
Characterization of healthy human iPSCs and iPSC-derived MSCs. **(A)** Identification of stem cell markers: ALP staining, fluorescent immunocytochemistry (ICC) of pluripotency markers TRA-1-60, SOX2, OCT4, SSEA4, and NANOG. Scale bar: 100 μm. **(B)** Embryoid body (EB) formation in suspension from aggregates of iPSCs. ICC of three germ layer markers: AFP, endoderm; α-SMA, mesoderm; and TUBB3, ectoderm. Scale bar: 50 μm. **(C)** Fluorescence-activated cell sorting of CD105+/CD45- MSCs. FITC, fluorescein isothiocyanate; PE, phycoerythrin. **(D)** The percentage of mesenchymal surface marker CD105+/CD45- cells in total cells differentiated from iPSCs originated from PBMCs and fibroblasts. Data are shown as mean + SEM. **(E)** and **(F)** Analyses of MSCs after sorting and expansion with flow cytometry for MSC negative markers (CD31, CD34, and CD45) in **(E)** and MSC positive markers (CD29, CD73, CD90, and CD105) in **(F)** labeled with different fluorochromes (V450, APC, PE, and FITC). Marker expression is presented as histograms (green). Unstained cells were used as controls (gray). Data are shown as mean + SEM. **(G)** Gene expression of MSC markers in transcripts per kilobase million (TPM). **(H)** Spindle-like morphology of iPSC-derived MSCs. Scale bar: 50 μm.

**Supplementary Figure 2.**
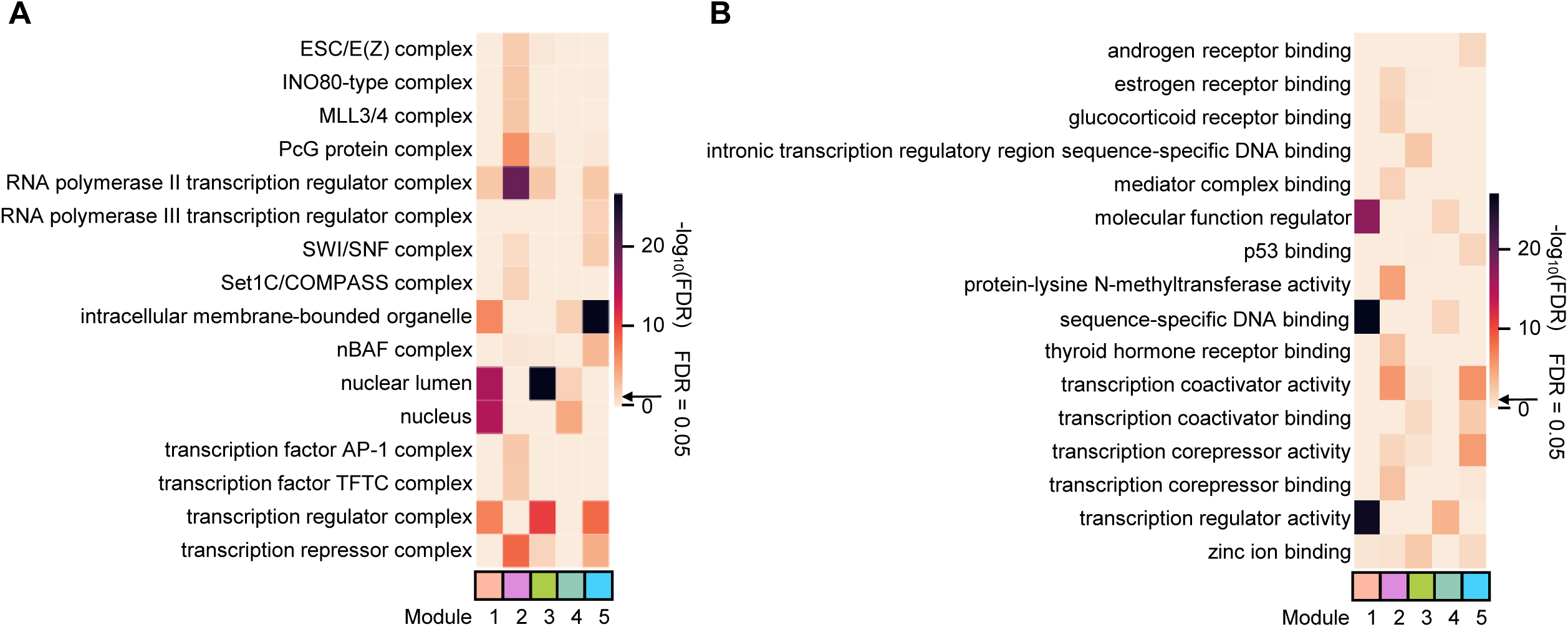
Gene Ontology (GO) enrichment of TF regulatory network modules. **(A)** Top enriched GO cellular component (CC) terms of each module. **(B)** Top enriched GO molecular function (MF) terms of each module.

**Supplementary Figure 3.**
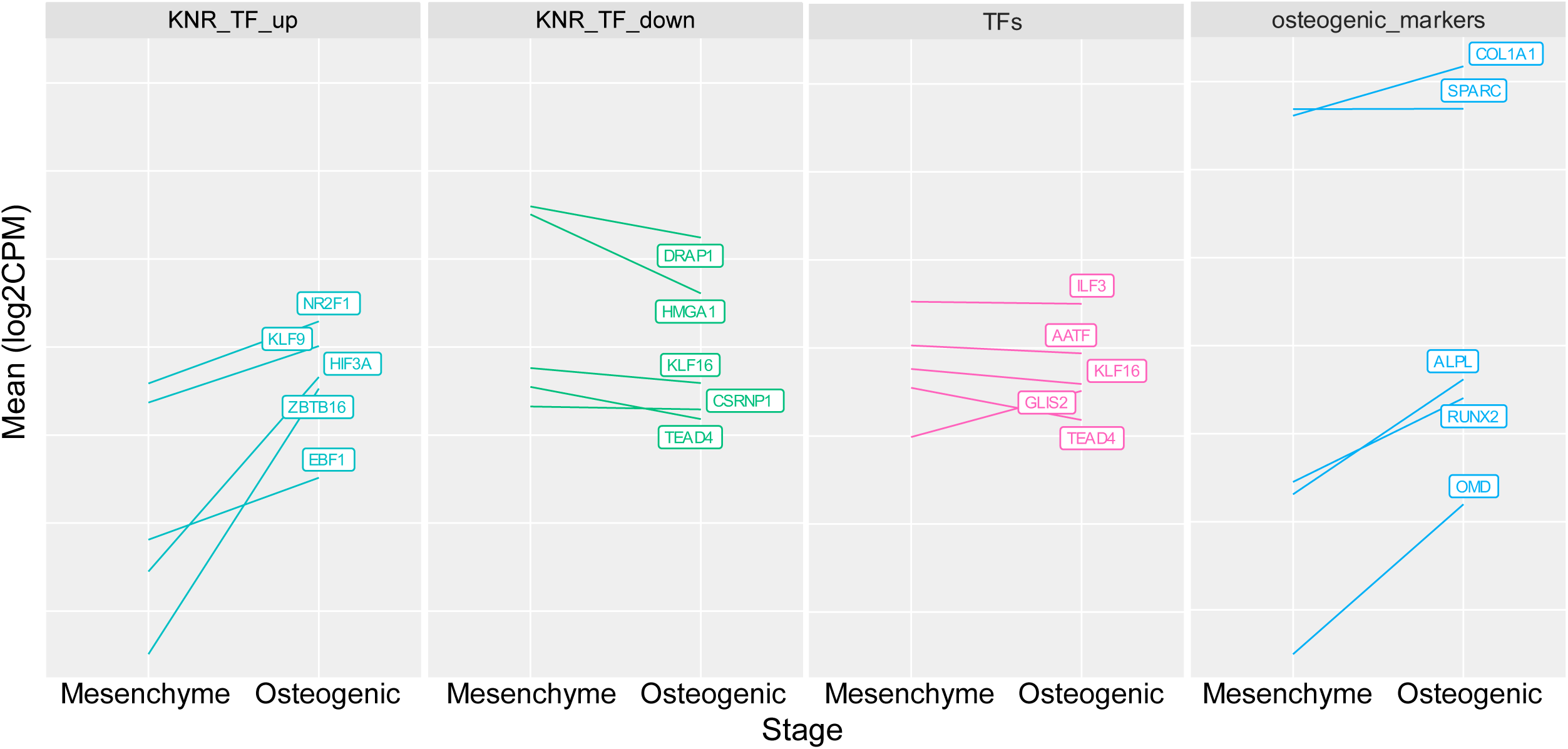
Gene expression pattern across osteogenic differentiation stages using pseudobulk single-cell RNA-seq data in Housman et al., 2022. KNR_TF_up: top five up-regulated KNR transcription factors; KNR_TF_down: top five down-regulated KNR transcription factors; TFs: top five transcription factors (TFs) based on TF regulatory network; osteogenic_markers: five known osteogenic markers.

**Supplementary Figure 4.**
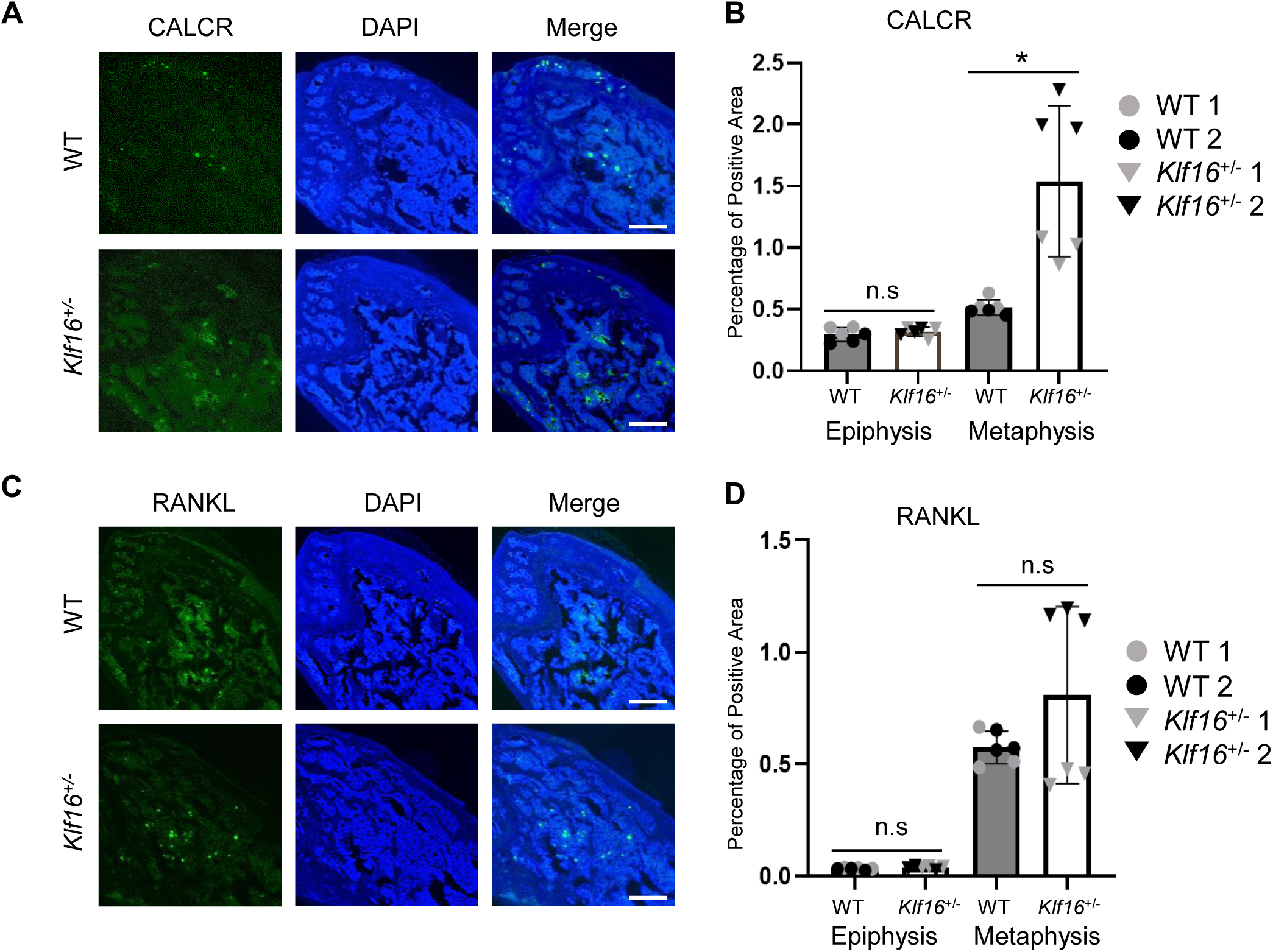
Expression of osteoclastogenesis markers in the femur bone of WT and *Klf16^+/-^* mice. **(A)** IHC for CALCR in the distal femur bone. The CALCR**-**positive areas in epiphysis and metaphysis of two *Klf16^+/-^* mice and two WT mice were quantified and shown as the percentage of CALCR-positive area (fluorescence-positive area/total bone area) in **(B)**. **(C)** IHC for RANKL in the distal femur bone. The percentage of RANKL-positive area were quantified and shown in **(D)**. For **(B)** and **(D)**, three replicate sections from each animal were analyzed. Data are presented as the mean ± SEM and visualized with GraphPad Prism 10. n.s: not significant. * p value < 0.05 by unpaired t-test. Scale bar: 500 µm.

**Supplementary Source Data for Figure 5, Supplementary Figure 1, and Supplementary Figure 4.**

**Supplementary Table 1**, related to Figure 1. Sample demographic metadata.

**Supplementary Table 2**, related to Figure 2. Differential gene expression between stages.

**Supplementary Table 3**, related to Figure 3. and Supplementary Figure 2. Transcription factor regulatory networks and gene ontology enrichments.

**Supplementary Table 4**, related to Figure 4. Significant MEGENA modules.

**Supplementary Table 5**, related to Supplementary Figure 3. Pseudobulk gene expression at osteogenic differentiation stages using single-cell RNA-seq data in Housman et al., 2022.

**Supplementary Table 6**, related to Supplementary Figure 3. Gene expression in five cell types at Day 21 of osteogenic differentiation using single-cell RNA-seq data in Housman et al., 2022.

**Supplementary Table 7**, related to Figure 5. Primers used for *Klf16* in vitro overexpression study.

